# Pipeline for Assessing Tumor Immune Status Using Superplex Immunostaining and Spatial Immune Interaction Analysis

**DOI:** 10.1101/2024.08.23.609368

**Authors:** Chaoxin Xiao, Ruihan Zhou, Qin Chen, Wanting Hou, Xiaoying Li, Yulin Wang, Lu Liu, Huanhuan Wang, Xiaohong Yao, Tongtong Xu, Fujun Cao, Banglei Yin, Ouying Yan, Lili Jiang, Wei Wang, Dan Cao, Chengjian Zhao

## Abstract

The characteristics of the tumor microenvironment (TME) are closely linked to tumor progression and treatment response. The TME comprises various cell types, their spatial distribution, cell-cell interactions, and their organization into cellular niches or neighborhoods. To capture this complexity, several spatial profiling technologies have been developed. However, challenges such as low throughput, high costs, and complicated data analysis have limited their widespread use in immune research. In this study, we introduce the Cyclic-multiplex TSA (CmTSA) staining platform, a high-throughput superplex staining technology based on tyramide signal amplification (TSA) immunostaining combined with an efficient fluorophore recycling method. The CmTSA platform allows for the labeling of 30-60 antigens across multiple parallel formalin-fixed paraffin-embedded (FFPE) slides. Furthermore, the automated CmTSA workflow requires only standard histological equipment and conventional immunohistochemistry (IHC) primary antibodies (Abs), significantly reducing costs. While the superplex images produced contain extensive multidimensional information, extracting the spatial features of the TME from raw pixel data can be challenging. To address this, we present a computer vision-based analysis pipeline, which begins with deep learning-based algorithms to segment individual cells and identify cell types based on defined annotation rules. It then evaluates the spatial distribution tendencies of each cell type, the interaction intensity between paired cells, and the multicellular functional niches. This comprehensive approach enables researchers to visualize and quantify the types, states, and levels of immune activities within the TME effectively, advancing tumor immunology research and precision immune medicine.

## INTRODUCTION

Tumor persistence, spread, and therapeutic response are influenced not only by the intrinsic properties of malignant cells but also by their interactions with various other cell types within the microenvironment. Immune cells, stromal cells, endothelial cells, adipocytes, and microbial cells can either promote or suppress tumors, depending on the context.^1,2^ Comprehensive understanding of cancer and effective treatment development require investigating the complex, multidirectional interactions and signaling networks between tumor cells and their surrounding microenvironment. In the past decade, single-cell RNA sequencing (scRNA-Seq) has been extensively used to map the gene expression profiles of tumor, immune, and stromal cells across various cancers.^3^ However, these techniques lack critical spatial information. Tumors function as ecosystems of heterogeneous cell types that dynamically interact and exchange information, and their spatial organization, including cell distribution and relationships with structures like blood vessels, influences disease progression and treatment outcomes.^4–6^ Traditional tissue architecture analysis using H&E stains, while crucial for identifying malignancy, often overlooks spatial interactions between tumor cells and other cell types. Therefore, molecular studies that preserve and reveal the spatial organization of tumors at the single-cell level are essential for understanding how intercellular communication shapes tumor growth and therapy response.

Profiling the spatial features of patients’ tumor ecosystems begins with identifying cell phenotypes in situ. The two mainstream technologies for this are spatial transcriptomics and spatial proteomics.^7,8^ Although spatial transcriptomics may offer more intracellular molecular information, its limited throughput makes it challenging to assess and compare spatial interactions among various cell types in the TME at the clinical cohort level TME.^9,10^ Additionally, mRNA-based detection methods face practical challenges for routine clinical samples due to the unstable nature of mRNA. Therefore, proteomics based on single-cell-resolved multiplex protein imaging is more suited for this purpose. It is compatible with routine clinical samples of large cohorts and can distinguish dozens of markers (typically more than 30), adequately characterizing cellular phenotypes and statuses within a tumor ecosystem. Spatial proteomics technologies can be grouped into two classes based on the readout signaling: mass spectrometry (IMC and MIBI-TOF) and fluorescence or chromogenic imaging (CODEX, CycIF, and multiplex IHC).^4,11–14^ Each method offers unique benefits and drawbacks in terms of resolution, multiplexing capacity, efficiency, and signal amplification. However, when applied to clinical samples, current technologies face significant challenges, including high background noise, low throughput, high costs, and incompatibility with conventional Abs and clinical samples. These issues limit their effectiveness and accessibility in clinical settings and hinder their application in research involving large cohorts.^15^ There is an urgent need to develop new methods to overcome these challenges, enhancing the reliability, efficiency, and affordability of TME spatial profiling processes.

Spatial proteomics technologies enable the visualization of dozens to hundreds of marker proteins across up to millions of cells within a tissue slide. This vast pixel dataset presents multidimensional information that characterizes the patient’s TME.^16^ It includes multiple layers of information such as single-cell phenotypes, single-cell morphology, composition of cells in the tissue, cell-cell interactions, and their arrangements into multicellular niches or larger anatomical/functional structures.^17^ However, extracting these features from the raw pixel data is not straightforward, and requires dedicated analysis methods, bridging the worlds of microscopic images, pathology analysis and computer science. Machine learning models (such as CellPose and StarDist) enable accurate cell segmentation and classification translate pixel-based high-plex imaging data into digitized, matrix-based spatial data.^18–20^ These computer-vision based methods could perform as accurately as humans when compared with manually segmented ground truth data. However, tumors often contain densely packed cells, some with irregular shapes (for example, fibroblasts), and z-dimension overlap that is not apparent in the x–y plane. Therefore, segmentation and cell type classification process must be conducted with care, and human visualization of images is generally needed to visually verify key results.

In addition, deriving biological and clinical insights from this novel data type remains an ongoing research challenge. To characterize the spatial interactions between cell types within tumors, many computational platforms have been introduced.^4,9,21,22^ Some cellular spatial analysis platforms focus on pairwise associations between cell types based on the frequency of spatial co-occurrence, spatial correlation, or nearest distances between individual cells. Other platforms use machine learning (e.g., K-means) or topic modeling to define tissue cellular niches with consistent cell type compositions.^4,17^ These platforms mainly focus on localized regions (small regions from TMA or specific ROI regions) rather than whole-mount pathological slides, lacking strategies to comprehensively understand the spatial characteristics of the patient’s immune microenvironment. Additionally, the behavior and spatial interactions of immune cells within the tumor immune microenvironment are complex but have inherent patterns. The subtypes, abundance, cell distribution tendencies, and spatial interactions with other immune cells within the microenvironment are all meaningful features that collectively define the immune function of the individual microenvironment.^17^ However, the commonly used spatial analyses are rather fragmented and lack logically interconnected systematic approaches, making it difficult to form an effective chain of spatial characteristic analysis of the immune microenvironment.^23^ This hampers the comprehensive and systematic evaluation of individual immune status.

We present a high-throughput, superplex staining method called Cyclic-multiplex TSA (CmTSA), capable of simultaneously targeting 30-60 markers across over 20 tissue slides. This method is compatible with conventional IHC Abs and instruments, producing highly amplified superplex images with minimal fluorescent background at a relatively low cost. Additionally, we introduce a computer vision AI-assisted cell segmentation model and a cell type annotation rule. These tools efficiently and accurately translate pixel-based superplex images into spatial matrix data, containing cell types and spatial positions of all segmented cells, and handling up to millions of cells. Based on this spatial matrix data, we provide a comprehensive workflow for profiling the spatial features of patients’ TME at the whole-mount slide level. In summary, we combine a new high-throughput and low-cost superplex staining technique, a computer vision AI-based digitization method, and a straightforward analysis pipeline for comprehensive profiling and quantifying the spatial characteristics and immune activities within patients’ TME at the whole-slide level.

## RESULTS

### Background fluorescence quenching of FFPE sections

Conventional FFPE sections exhibit significant autofluorescence. This background fluorescence arises from collagen fibers, elastin, red blood cells within the tissue, and aldehyde compounds (formaldehyde and formalin) used during tissue fixation.^24^ The emission spectra of these autofluorescent molecules overlap substantially with the excitation spectra of commonly used fluorescent labeling molecules, thereby significantly interfering with the detection specificity of labeled fluorescence (Fig. 1a). To overcome this problem, we developed an effective approach that combines photobleaching and chemical quenching for eliminating tissue background fluorescence. In this method, deparaffinized sections were uniformly irradiated with a high intensity refrigerated LED light source (600W, with continuous emitting spectrum) while submerged in a solution containing 4% hydrogen peroxide. The intense LED-light excites various potential background fluorescent molecules within the section, while the 4% hydrogen peroxide in the solution photolyzes to produce hydroxyl radicals (·OH). These hydroxyl radicals (·OH) undergo redox reactions with the excited fluorescent molecules, altering their structure, which changes their energy transitions and results in fluorescence quenching (Fig. 1b). Test results indicate that a 8-minute quenching treatment can effectively eliminates more than 90% background fluorescence from paraffin sections without affecting the detecting ability for various antigens (Fig. 1c, d, f, Supplementary Fig. 1). Further evaluation results indicate that this method is compatible both to conventional immunofluorescence (IF) staining and covalent TSA staining (Fig. 1e), providing a widely useful way to efficiently eliminate background fluorescence and dramatically enhance the staining Signal-to-Noise Ratio (SNR) during various fluorescence labeling process (Fig. 1f, Supplementary Fig. 1).

**Fig 1.**
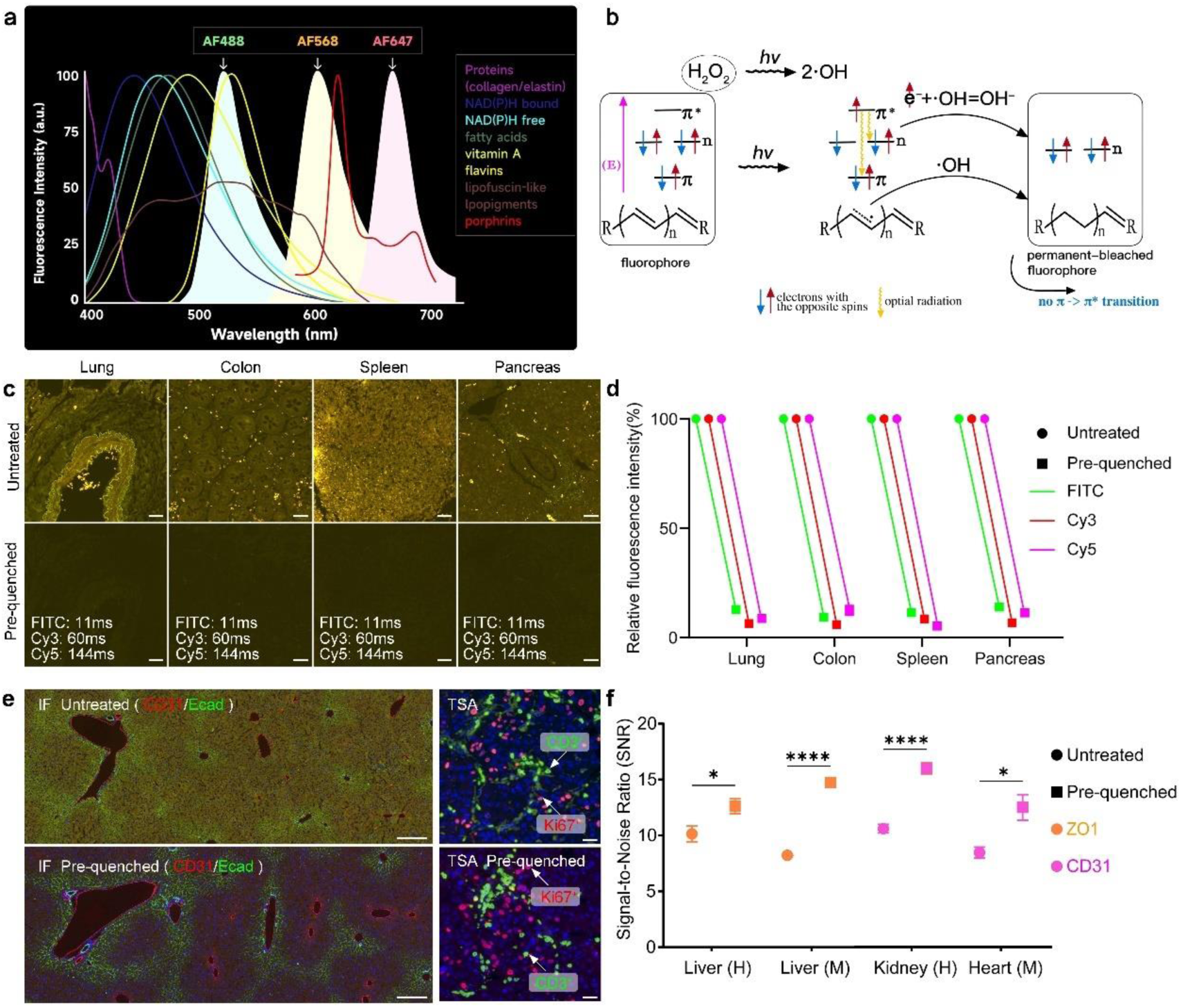
Background fluorescence quenching of FFPE sections. (a) The emission spectra of autofluorescent molecules overlap substantially with the excitation spectra of commonly used fluorescent labeling molecules. (b) The principle of background self-fluorescence quenching. (c) Background fluorescence quenching techniques can significantly reduce the background fluorescence in various human FFPE sections. The same exposure time is used for each imaging channel on every slice. Scale bar, 50 μm. (d) The quenching efficiency of background fluorescence in various human tissues. (e) Whether using IF staining (in mouse liver) or TSA staining (in human tonsil), background fluorescence quenching can significantly improve the Signal-to-Noise Ratio (SNR) of staining images. Scale bar of IF staining, 100 μm. Scale bar of TSA staining, 20 μm. (f) The SNR statistics of different antibody markers after background fluorescence quenching. Data shown as Mean ± SEM. *: p ≤ 0.05; ****: p ≤ 0.0001. Based on Unpaired Student’s t-test.

### Gentle antibody stripping is necessary for sequential immunostaining with scores of iterations

Similar to IF labeling, most superplex protein labeling methods are also based on the specific binding of antigens and antibodies (Abs). The main strategy to achieve superplex labeling currently is by directly using panels of pre-labeled primary Abs (from dozens to hundreds). Based on the different read-out methods, these specifically designed Abs are pre-labeled with various tags (e.g., Codex using DNA tags, IMC using rare metal tags, and CycIF technology using fluorescent molecule tags).^4,11,25^ While these superplex labeling technologies are increasingly applied in the studies of tissue microenvironments, particularly immune microenvironments, several concerns have been raised. Firstly, the specificity and labeling performance of these modified Abs may vary from traditional primary Abs validated by IHC, and researchers cannot independently verify them before use. Secondly, the commercial availability of pre-labeled primary Abs is limited, often failing to meet the specific needs of individual studies and thus restricting research design. Lastly, the direct labeling approach can result in low signal intensity, high background noise, and reduced resolution in fluorescence or mass spectrometry imaging.

These issues are particularly evident for low-abundance antigens, reducing image quality and affecting the reliability of final analysis results.

To replace the pre-labeled Ab with conventional Ab, indirect immunolabeling methods are required. However, during multiplex labeling, potential cross-binding of 1^st^ Abs to the fluorophore-labeled 2^nd^ Abs has to be considered and avoided, which limits the co-labeling capability of IF method to 2-3 targets. In contrast, during TSA staining, fluorophore was not attached on 2^nd^ Abs, but covalently bind to antigens via the Tyramide-HRP reaction, which allowing Ab stripping without affecting specific fluorescent labeling. Thus, continuous Ab stripping enables independent sequential multiplex labeling, even using the 1^st^ Abs originating from the same species. However, traditional Ab stripping is mainly conducted through microwave boiling (at temperatures above 95°C in acidic or basic buffers), which is simple to operate but causes damage such as cell loss and tissue detachment, particularly evident in tissues or regions with low cell density.^26^ In addition, continuous high-temperature boiling may potentially affect subsequent antigen epitopes and labeling efficiency.

To address this issue, our research group summarized and evaluated various potential Ab stripping reagents (such as solution salt concentration, pH, and surfactants). We found that using the reducing agent beta-mercaptoethanol to depolymerize Ab disulfide bonds is the most efficient and least damaging method for homologous Ab stripping (Fig. 2a, Supplementary Fig. 2). While using a low PH buffer is the best strategy to avoid the cross labeling when encountering a heterologous 1^st^ Ab (Fig. 2a). Test results demonstrate that our Ab stripping reagent effectively removes both homologous and heterologous antibodies, allowing for successful multiplex TSA staining in both FFPE sections and cultured cells (Fig. 2b, c). Importantly, the morphology of synapses in cultured neurons remained intact, indicating that this Ab stripping method is gentle and preserves tissue and cellular structures (Fig. 2c).

**Fig 2.**
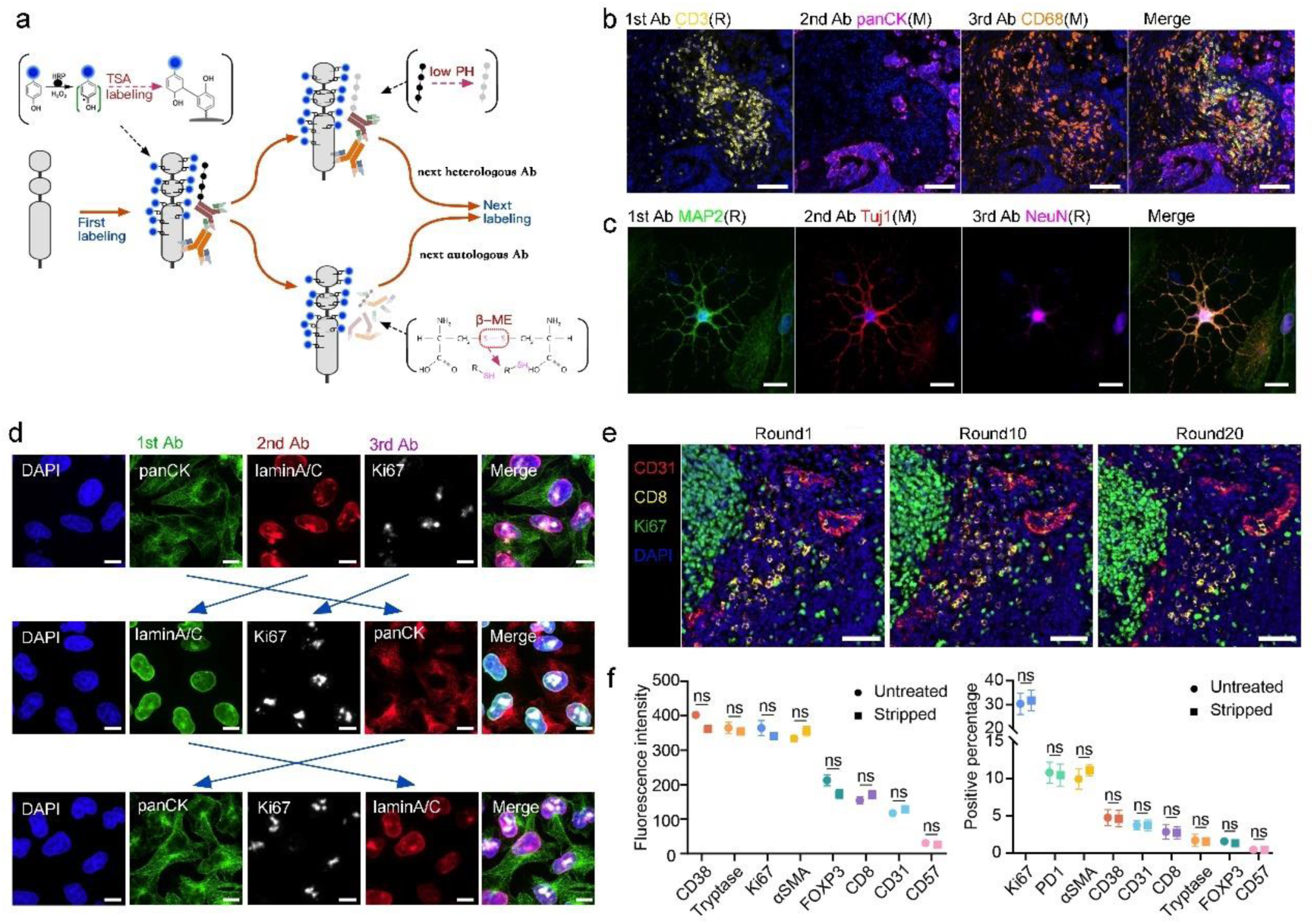
Gentle antibody stripping is necessary for sequential immunostaining with scores of iterations. (a) Schematic depicting TSA staining and antibody stripping. (b) Representative images of multiplex TSA staining of human lung cancer FFPE sections. (R), the antibody source is rabbit. (M), the antibody source is mouse. Scale bar, 50 μm. (c) Representative images of multiplex TSA staining of cultured neuron cells. Scale bar, 20μm. (d) Altering the order of antibody labeling in the multiplex TSA staining process for HeLa cultured cells produces consistent results. Scale bar, 20 μm. (e) Representative images of multiplex TSA staining after 0, 10, and 20 consecutive rounds of antibody stripping on three consecutive sections of human tonsil tissue. Scale bar, 50 μm. (f) Fluorescence intensity and positive percentage statistics for various antibody markers on human tonsil FFPE section after 20 consecutive rounds of antibody stripping. Data shown as Mean ± SEM. ns, not significant, based on Unpaired Student’s t-test.

To assess the impact of the Ab stripping reagent on protein integrity, location, and epitope exposure, we varied the labeling order during multiplex TSA staining. Our results were consistent across all staining sequences (Fig. 2d). Additionally, we conducted 20 consecutive rounds of Ab stripping and found that antigen epitopes and fluorescent labeling capabilities remained unaffected (Fig. 2e, f). These findings indicate that the Ab stripping method is efficient and gentle, preserving epitope features, cell morphology, and tissue structure during multiple rounds of sequential TSA staining.

### Highly efficient cyclic reuse of fluorophores

Achieving multiple fluorescence labeling requires technologies or strategies that can effectively distinguish different fluorescence signals. Currently, there are two main methods in use: spectral unmixing and cyclic imaging. The former utilizes the unique emission spectra characteristics of each staining molecule to distinguish different labeled fluorescence signals through optical separation and complex algorithms. The latter uses multiple cycles of fluorescence combinations (such as FITC, Cy3, and Cy5 fluorescence combinations). Spectral unmixing only requires a single imaging session, but it can currently only separate up to 9 fluorescent colors in section image data.^27,28^ On the other hand, cyclic fluorescence imaging requires multiple imaging sessions and image registration, but theoretically, its ability to distinguish the number of fluorescence signals is unlimited. In our previous studies on background fluorescence quenching in FFPE sections, we established an efficient background fluorescence quenching method. We found that by altering the excitation spectra, this method could achieve differential quenching of fluorescent molecules (e.g., quenching AF488, AF561, FITC, Cy3, and Cy5 labeled fluorescent molecules, while not quenching the DAPI-labeled nuclear fluorescence signal). Tests demonstrated that even for FFPE sections labeled with fluorescence using the TSA method (where fluorescence intensity is 5-100 times higher than traditional IF staining) (Fig. 3a, Supplementary Fig3), our method achieved nearly 100% quenching of labeled fluorescence signals within 10 minutes (Fig. 3b).

**Fig 3.**
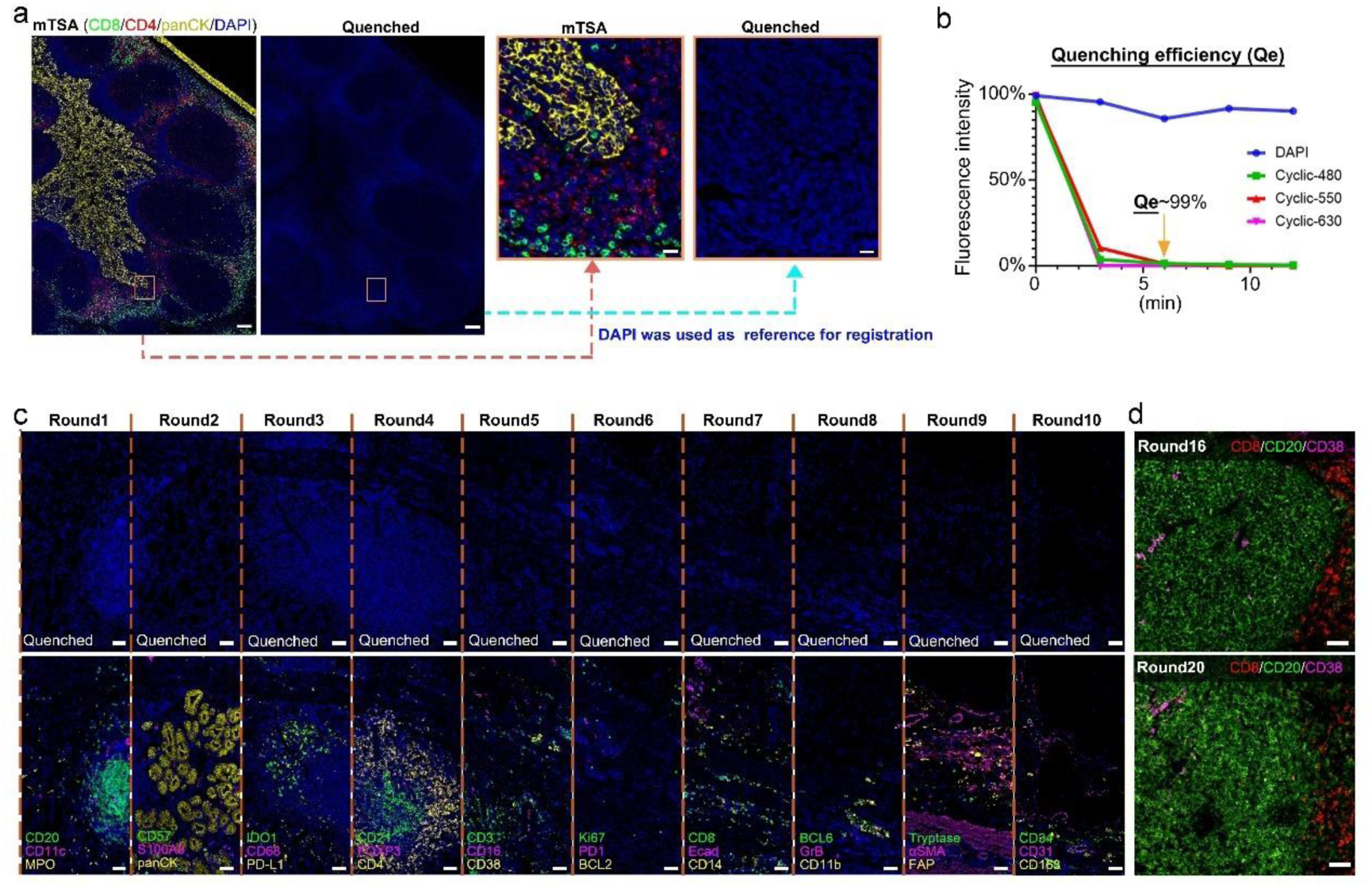
Highly efficient cyclic reuse of fluorophores. (a) Representative images showing differential quenching of fluorescent molecules marking CD8, CD4, and panCK through mTSA on a human tonsil FFPE section. Scale bar, 200 μm. Zoom in scale bar, 20 μm. (b) The quenching efficiency of different fluorescent molecules. (c) Representative images of 10 rounds CmTSA staining, performed after 10 rounds of continuous fluorescence quenching on a human gastric FFPE section. Scale bar, 50 μm. (d) Representative images of two human tonsil FFPE sections stained with the same antibody after 16 and 20 rounds of continuous fluorescence quenching. Scale bar, 50 μm.

Further tests indicated that for up to 20 rounds quenching process did not significantly affect following antigen detection (Fig. 3c, d).

### Computer vision-based registration of serial images

Combing the multiplex TSA and the cyclic imaging, we developed a Cyclic multiplex TSA super staining method (CmTSA), which can yield a series of images that are collected separately. To properly process these data and correctly assign markers to individual cells, it is essential that all the images from the same FFPE section be accurately aligned. While various registration algorithms have been reported, ^29,30^ we sought a method that could be applied to align large datasets (each whole clinical tumor section could contain 0.1-1 million cells). For each cycle of staining, DAPI was not quenched and served as coordinates for image registration (Fig. 4a). For these reasons, we developed a workflow using Scale-Invariant Feature Transform (SIFT) keys, a computer vision algorithm used for object recognition proposed by David Lowe.^31,32^ SIFT algorithm extracts local feature points from an image and enable registration of serial images.

**Fig 4.**
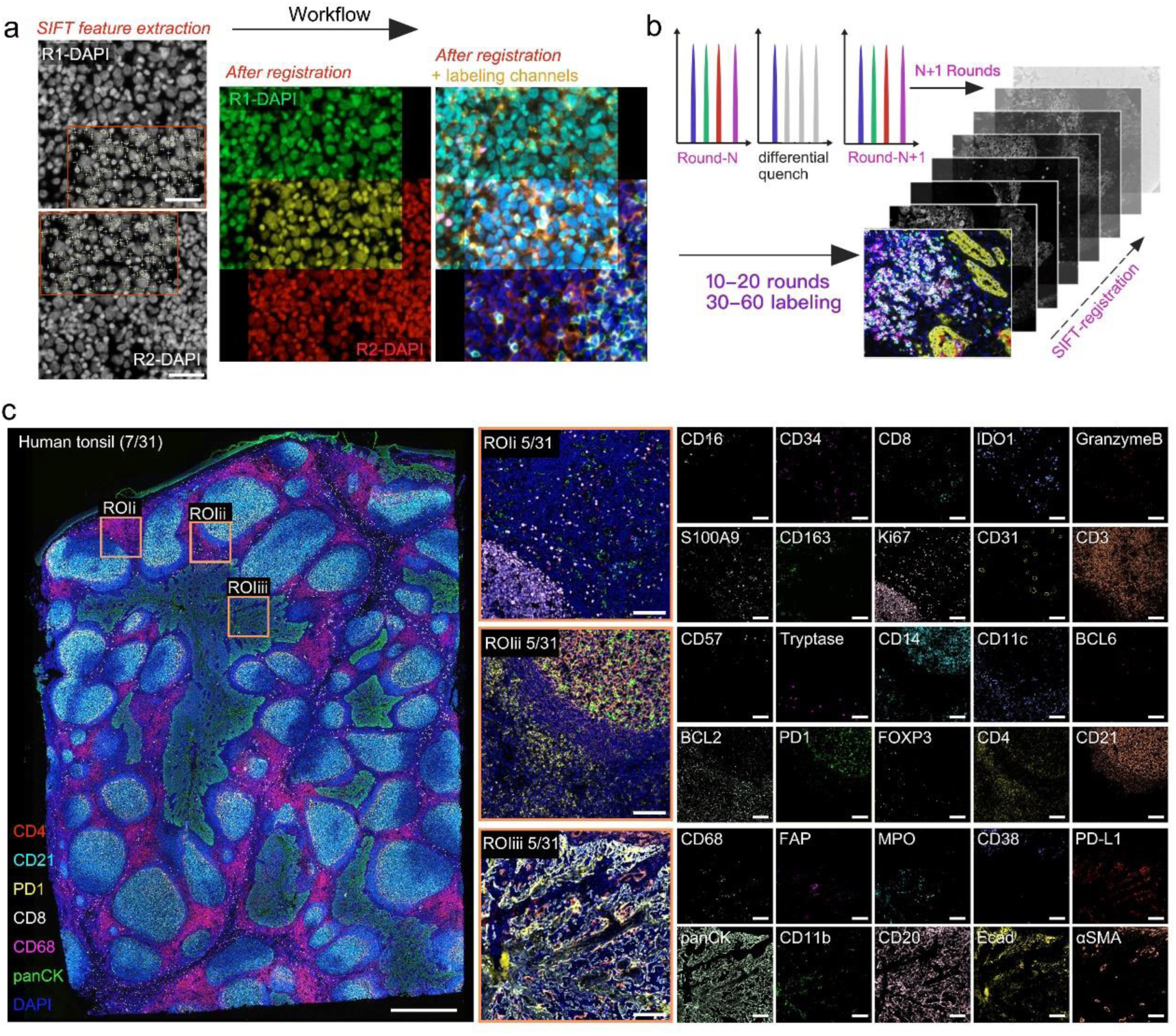
Computer vision-based registration of serial images. (a) The workflow of image registration based on SIFT feature extraction. Scale bar, 20 μm. (b) Schematic depicting cyclic-imaging and serial images registration based on SIFT. (c) The registered image of 30 Abs marked on a human tonsil section, which underwent 10 rounds of CmTSA staining. Wholesilde scale bar, 1 mm. ROI scale bar, 100 μm.

SIFT features are robust to various transformations: they are invariant to different scales, effectively handling image scaling; invariant to image rotation; partially invariant to changes in illumination; and invariant to a certain extent of affine transformations, such as image skewing and stretching. These excellent features are particularly suitable for accurately aligning serial images that derived from CmTSA. Due to our differential quenching method, DAPI signaling was saved and utilized as the coordinates/common features for SIFT to align the serial multichannel images (Fig. 4b). To test the fidelity of this method, a 10-rounds CmTSA staining experiment was performed on human tonsil sections labeled with DAPI and other 30 Abs (Fig. 4c). For this experiment, DAPI of the 1st round image was selected as the coordinates used for the registration of other 9 serial multichannel images. The alignment results showed perfect pixel-to-pixel matching of fluorophore signaling to the first DAPI (Supplementary Fig. 4). In addition, the method allows alignment of large datasets (image from whole slide scanning, containing up to 1 million cells).

### Cell segmentation and cell type defining from superplex images

Using our CmTSA method, up to 20 rounds of multiplex staining can be performed before significant damage to antigens and tissue structure occurs, generating a pathological superplex image with up to 1 million cells (typical whole tumor FFPE slide) and 60 signaling channels. The vast and complex information in these images is too extensive for traditional analysis methods by researchers or pathologists through human visual inspection. Thus, processing and analyzing this superplex image data requires computer vision AI. The first essential task is accurate cell segmentation and cell type annotation. We tested and applied multiple deep learning models, such as StarDist, Cellpose, and TissueNET, for this purpose. StarDist represents cells using star-convex polygons, approximating each cell’s shape with multiple rays extending from a central point to delineate the cell’s boundary.^31^ This model is particularly suitable for segmenting morphologically diverse cells in complex tumor microenvironments, where cells are densely arranged and frequently overlapping. We used a StarDist pretrained model to segment a human cervical cancer FFPE section staining image and achieved excellent results in QuPath (Fig. 5a).^33^ Following cell segmentation, cell type annotation involves two major steps, which we refer to as the cell gating strategy: Firstly, determine the expression level of cellular markers in individual cells, get the single positive marked cell; Secondly, design an annotation rule to gate cellular types or subtypes. The expression patterns of cell markers, revealed through fluorescent signals, vary. Several key issues must be carefully considered when applying this methodology. Some markers are expressed on the cell membrane (e.g., CD8 and CD4), some in the cytoplasm (e.g., panCK and CD68), and others in the nucleus (e.g., FOXP3 and Ki67). Different extraction methods must be carefully used basing on the specific expression patterns of individual markers (Fig. 5b). Additionally, since certain antigen markers are shared by multiple cell types and some cell types require multiple markers to define, an annotation rule is essential for ensuring consistent, accurate, and automated classification of cellular subtypes. For example, we defined Treg cells by FOXP3 and CD4 double positive marked cells (Fig. 5c). To benchmark how the cell gating strategy performs, we analyzed a superplex image of human tonsil stained by CmTSA (Fig. 5d, e). A panel of lineage-defining markers and cell type annotation rule (based on typical lineage markers, expected population abundance, and staining quality supervision) was used to define the cell types within the superplex image (Supplementary Fig. 5a). We observed strong agreement between the ground-truth multiplex images and the predicted results (Supplementary Fig. 5b), demonstrating that an appropriate cell gating strategy enables accurate classification of the diverse cells present in the superplex images.

**Fig 5.**
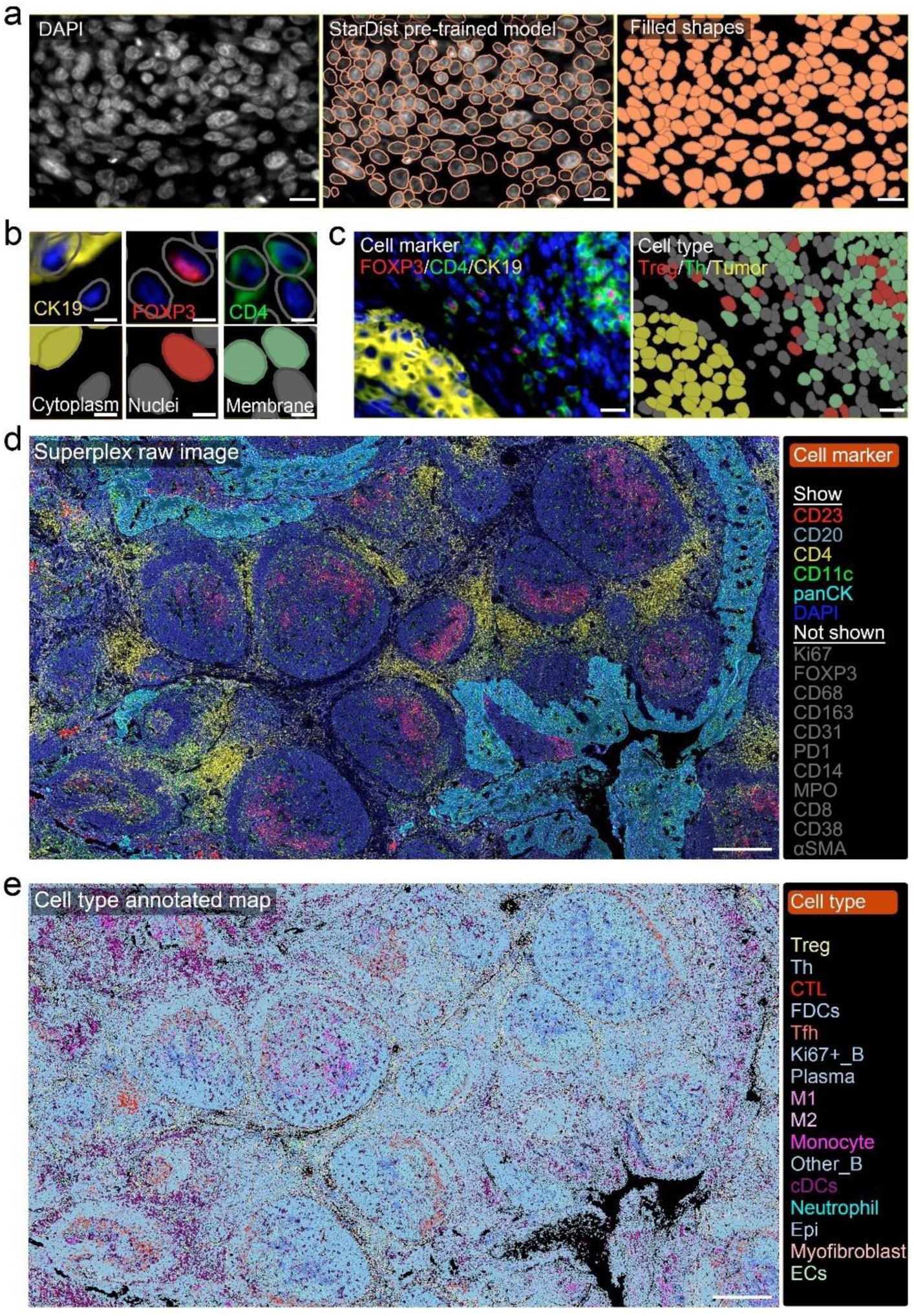
Cell segmentation and cell type defining from superplex images. (a) Cell segmentation of stained images from human cervical cancer FFPE sections using the StarDist pre-trained model. Scale bar, 20 μm. (b) Different extraction methods used to determine the expression levels of individual Abs labeled on human cervical cancer FFPE section. The gary color masked cell represent negative cell. Scale bar, 5 μm. (c) Cell type defining of a mTSA image based on positive marker results. Scale bar, 20 μm. (d) Superplex image(16 antibodys marker) of a human tonsil FFPE section stained by CmTSA. Scale bar, 500 μm. (e) Cell type defining after cell type annotation rule. Scale bar, 500 μm.

### Evaluation of spatial distribution tendencies for individual types of immune cells

In the TME, immune cells are unevenly distributed due to several factors. Tumor heterogeneity creates regions with varying levels of oxygen, nutrients, and growth factors, affecting immune cell infiltration.^2,34^ Chemokine gradients attract or repel specific immune cells, leading to clusters. Immunosuppressive zones with regulatory T cells and myeloid-derived suppressor cells (MDSCs) inhibit effector immune cell activity.^35,36^ The dense extracellular matrix(ECM) and abnormal vasculature hinder immune cell migration, causing them to accumulate in accessible areas.^37^ Additionally, varying metabolic conditions, such as hypoxia and acidic pH, create hostile environments for certain immune cells.^38^ These factors lead to specific distribution patterns of immune cells within the TME, potentially reflecting their roles in the immune response, such as targeting tumor cells, modulating inflammation, or suppressing immune activity. ^6,39^ These distributing patterns also reveal the overall immune landscape of the tumor, including areas of immune activation, suppression, or evasion.^40,41^

To describe the spatial distribution features of cells in the TME, we introduced the degree of aggregation and the Infiltration Score (Fig. 6a). Using the CmTSA method and a computer vision-based AI model, we annotated cell types and their spatial locations within the TME, creating a spatial matrix of all annotated cells. This matrix serves as the foundation for analyzing the spatial distribution features of immune cells. Using data from a cervical cancer patient (CC_P1) as an example (Fig. 6b), we evaluated the distribution tendency of immune cells by calculating the ratio of the average same-cell distance to the maximum same-cell distance (see MATERIALS AND METHODS). Permutation analysis indicates that the distribution of the different immune cells within the TME is neither random nor uniform (Fig. 6c), with all different kinds of immune cells showing a tendency to cluster to varying degrees. This clustering tendency of immune cells can be indicative of their coordinated efforts to exert their effects, such as targeting tumor cells or modulating the immune response.

**Fig 6.**
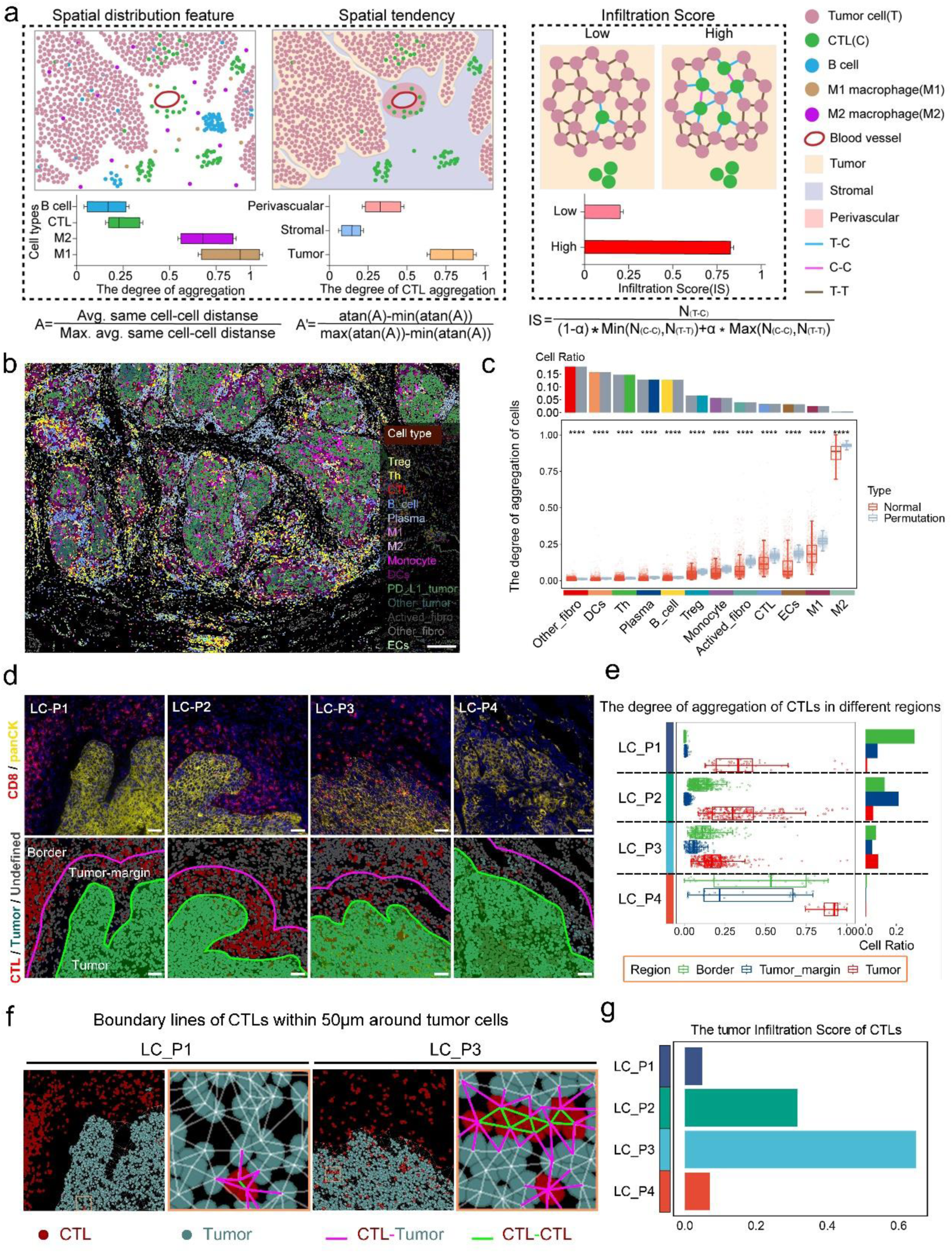
Evaluation of spatial distribution tendencies for individual types of immune cells. (a) A diagram illustrating the spatial distribution tendencies of cells within theTME. (b) The cell type defined mask rendered image of cervical cancer patient 1(CC_P1). Scale bar, 200 μm. (c) The spatial aggregation analysis of cells, excluding tumor cells, between the normal group and the permutation group was conducted using an Unpaired Student’s t-test. **** indicates P < 0.0001. (d) Representative images of four lung squamous cell carcinoma patients whose CTLs are defined in different areas. Scale bar, 50 μm. (e) The degree of aggregation of CTLs in three different areas within the four patients. Similarly, values closer to 0 also indicate stronger aggregation. (f) Representative images showing the boundary lines of CTLs within 50μm of tumor cells. Scale bar, 50 μm. (g) The tumor Infiltration Score of CTLs of four lung squamous cell carcinoma patients.

In addition, spatial aggregation analysis of immune cells from different TME regions (tumor or stroma) revealed varying degrees of spatial clustering among the same type of immune cells, particularly for Treg, Th, M1, plasma cells, B cells, monocytes and fibroblasts (Supplementary Fig. 6a-c). The varying degrees of spatial distribution tendencies of individual immune cell type in different regions may reflect their heterogenous function/role in responding to localized signals (either activation or suppression) within the TME.

To correlate the spatial distribution patterns of immune cells with clinical outcomes, we evaluated the spatial distribution tendencies of immune cells in different patients who received the same neoadjuvant anti-PD-L1 immunotherapy (see MATERIALS AND METHODS) (Fig. 6d). Given the critical role of cytotoxic T cells (CTLs) as mediators of tumor killing during anti-PD-L1 immunotherapy, our analysis focused on CTLs. Evaluation of surgical specimens revealed significant variability in CTL frequency and aggregation within different subregions (Tumor, Tumor-margin, and Border) among patients, despite all having a pathological partial response (pPR) (Fig. 6e). To quantify the degree of CTL aggregation within the tumor mass, we introduced an infiltration score, which highlighted the differences between patients and provided a more intuitive measure (Fig. 6f, g, Supplementary Fig. 6d). These findings demonstrate dramatic variability in the spatial distribution tendencies of CTLs within the remaining tumor tissue after neoadjuvant anti-PD-L1 immunotherapy, even among patients with the same pathological response. This variability suggests the possibility of differing prognostic outcomes for these patients. ^42–44^

### Detection and quantification of the spatial interaction between paired immune cells

In TME, immune cells often perform their functions by interacting with other immune cells or other local cells (Fig. 7a). For example, dendritic cells (DCs) present antigens to T cells through MHC molecules, initiating adaptive immune responses; follicular helper T cells (Tfh) interact with B cells to promote Ab production, indicating the effectiveness of humoral immune responses; regulatory T cells (Tregs) suppress the activity of effector T cells to maintain immune homeostasis and prevent autoimmune reactions; natural killer (NK) cells release IFN-γ to activate macrophages, enhancing their killing functions; tumor-associated macrophages (TAMs) interact with tumor cells by secreting immunosuppressive factors and promoting tumor invasion.^45–49^ Thus, identifying and quantifying paired cell interactions within the TME will undoubtedly enhance our understanding of the immune status of the TME in individual patients or patient cohorts.

**Fig 7.**
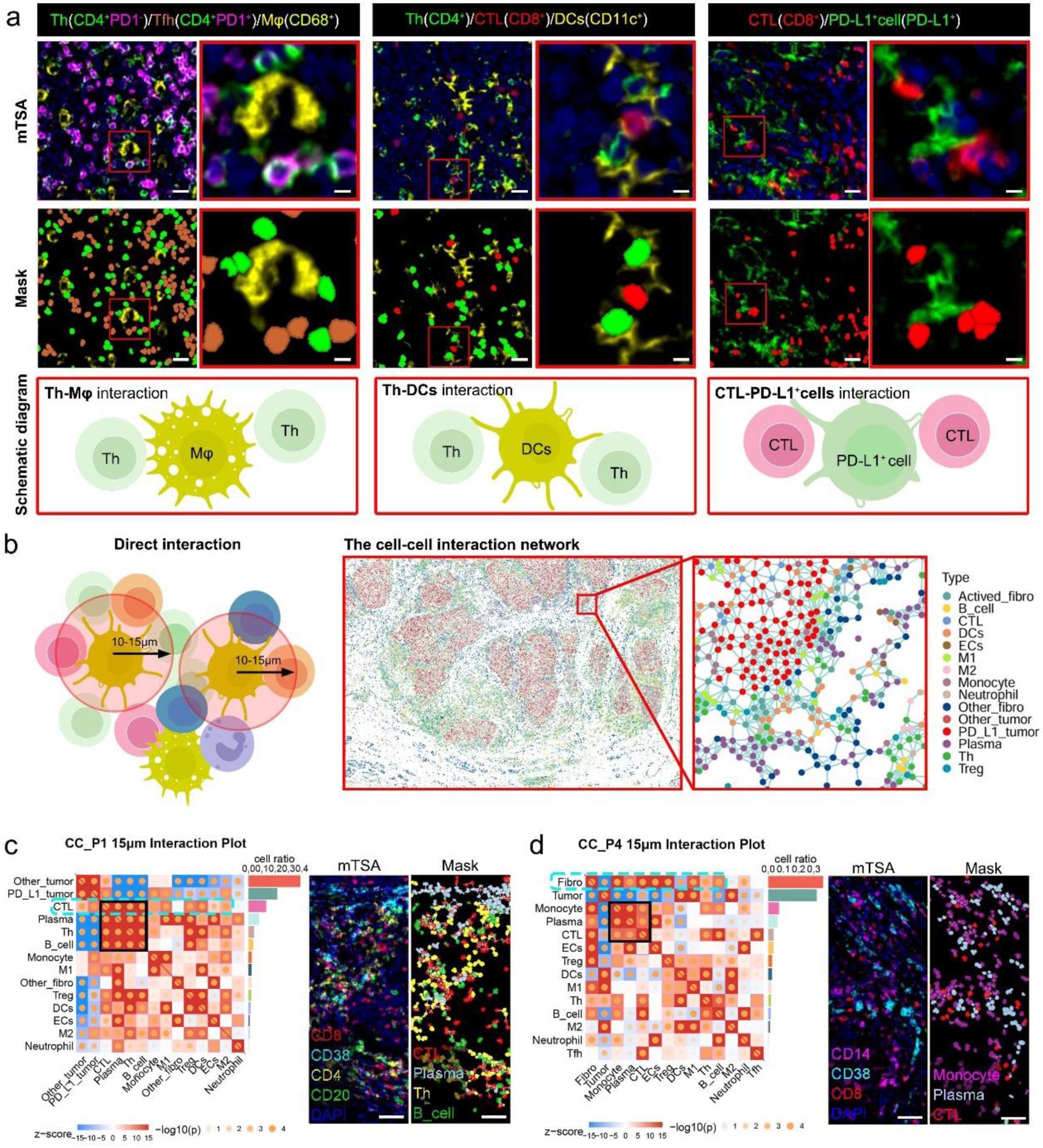
Detection and quantification of the spatial interaction between paired immune cells. (a) Representative images of cell-cell interaction. Left: the interaction of Th and Mφ in human tonsil. Middle: the interaction of Th and DCs in human tonsil. Right: the interaction of CTL and PD-L1^+^cell in human cervical cancer. Scale bar, 20 μm. Zoom in scale bar, 5 μm. (b) Diagrams illustrating direct interactions between cells and the overall cell-cell interaction network. (c,d) Heatmap of paired cell interactions within a 15 μm radius and partial representative images corresponding to CC_P1 (c) and CC_P4 (d). Scale bar, 50 μm. Colors in the heatmap represent Z-score, with deep blue indicating negative correlations and deep red indicating positive correlations. Circles represent -log10(p), used to indicate the significance of interactions.

Combining the CmTSA method with a computer vision-based AI model, we created a spatial matrix of the TME with annotated cell types and cell coordinates (Fig. 6a). Based on this information, we constructed a cell interaction network (see Material and Methods), where each cell is represented as a node and edges are created between nodes (cells) within a certain distance, indicating potential interaction (Fig. 7b). This interaction network can cover the entire region of interest (ROI) or even an entire slide, containing up to millions of nodes/cells.

Using spatial data from two cervical cancer patients, we measured the level of direct interaction between each cell type and all other cell types in the TME (Figure 7c, d). Interestingly, we found that the most active cell type differed between the patients: CTLs were the most active in CC_P1(Fig. 7c), while fibroblasts were the most active in the CC_P4(Fig. 7d). Additionally, by identifying groups of cells that interact more frequently with each other than with the rest of the network, we predicted potential functional cellular clusters within the TME. A CTL-Plasma-Th-B cell cluster was found in CC_P1(Fig. 7c), while a CTL-Plasma-Monocyte cluster was identified in CC_P2 (Fig. 7d), indicating different immune statuses between these two patients.

### Prediction and quantification of multicellular immune niches at the whole-slide scale

In addition to cell-to-cell interactions, the tumor immune microenvironment hosts higher spatial arrangements of multiple cell types. Different terms have been used for describing the local multi-cellular structures (neighborhood, community and milieu. et al.).^50–52^ Local multi-cellular structures are established and maintained through cell-cell talks mediated both by ligand-receptor binding (e.g., MHC-II/MHC-I to TCR, CD80/CD86 to CD28/CTLA4, and PD-L1 to PD-1) and cytokine secretion (e.g., IL-12, IFN-γ, IL-10, and TGF-β) ^53–56^ (Fig. 8a). Typically, ligand-receptor interactions occur between paired cells, paracrine signaling can extend to several adjacent cells, but its range could be significantly influenced by the uptake of surrounding targeting cells or competitors, such as Treg consuming IL-2.^3,57^ Considering the inherent regulation of multi-cellular interactions in immune activity, describing and quantifying the multi-cellular immune niches (INs) within the TME may be more accurate for the manifestation of the functional immune units and immune status of a patient.^58–60^

**Fig 8.**
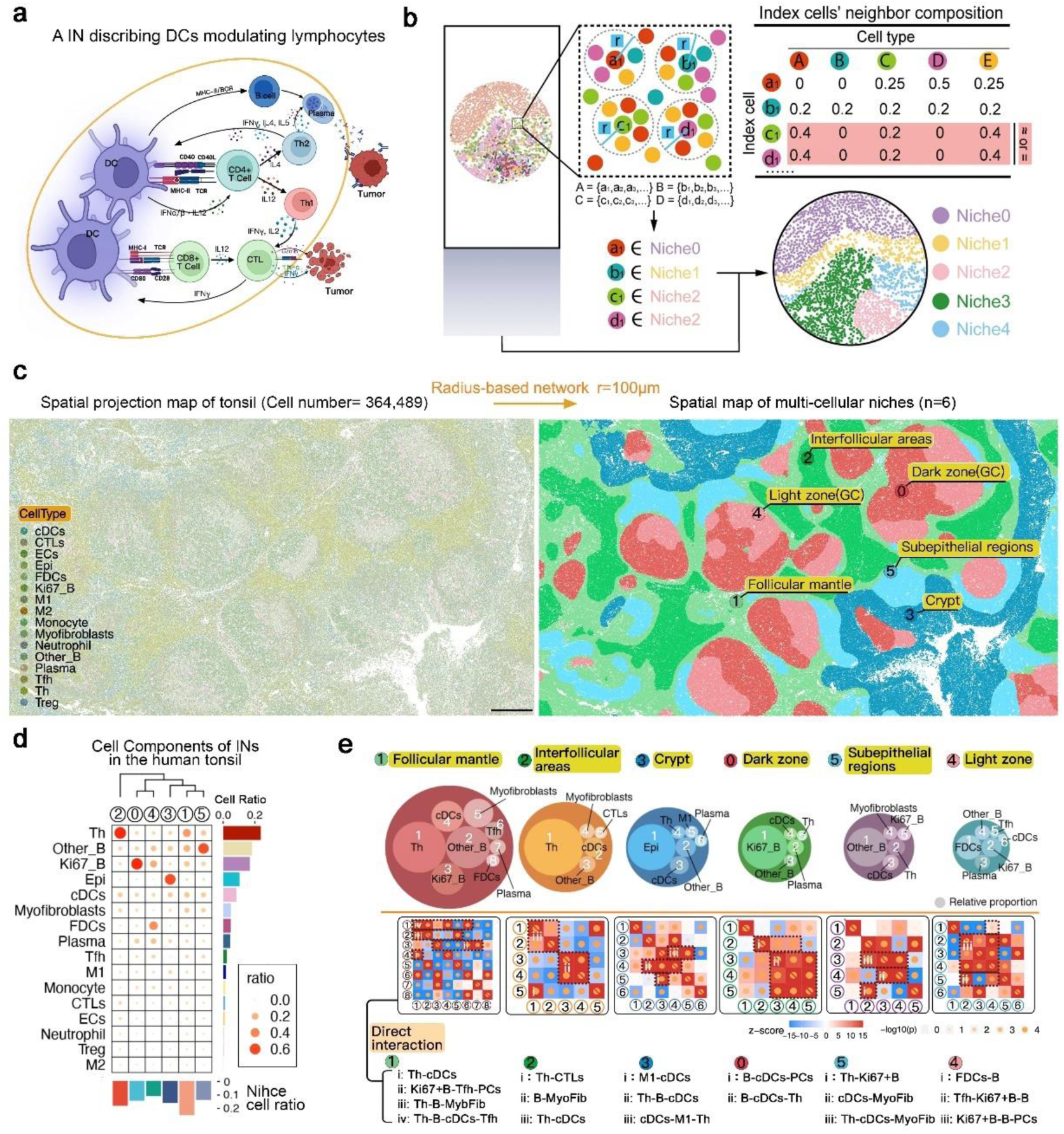
Prediction and quantification of multicellular immune niches at the whole-slide scale. (a) Diagrams illustrating cell-cell communication mediated by ligand-receptor interactions. (b) Diagrams illustrating the identification of INs using a radius-based network. (c) Spatial projection map and spatial map of multicellular niches in the tonsil, following CmTSA multiplex staining and cell annotation. Scale bar, 500 μm. (d) Bubble plot of the cellular composition of the six niches in the tonsil sample, with color and size representing the ratio of cells in each niche. (e) Top: Composition of the top 90% of cells in six tonsil niches. Large circles show niches, with smaller circles representing different cell types. Circle size indicates cell ratio; Middle: Heatmap of direct interactions between the top 90% of cell types in each niche. The meaning of the heatmap is the same as in Fig. 7c, d. Black dashed lines highlight significant direct interactions; Bottom: Labels indicating the categories of direct interactions between specific cell types within each niche.

The formation of INs is mainly determined by various spatial interactions between neighboring cells through ligand-receptor binding or cytokine secretion. To classify each cell within the TME into INs, we constructed a radius-based network. This approach was chosen because spatial interactions between neighboring cells are typically constrained by distance rather than a fixed number of neighbors. We determined each cell’s neighbors using a fixed radius, instead of using a fixed number of neighbors. Then, we applied the K-means clustering algorithm to categorize these cells and identify distinct INs (Fig. 8b, Material and Methods). This approach is more accurate in the TME, where cell densities vary and interactions are distance-limited (e.g., ligand-receptor binding and cytokine secretion). The radius value represents the true range of cell interaction, which can range from about 4 to 20 cells or 40-200 μm in diameter, but it highly depends on the type of surrounding cells and the features of local microenvironment.

To validate the feasibility of using the radius method for INs prediction, we tested a normal human tonsil. The tonsil, as a well-defined human immune organ, provides an excellent positive control for validating the ability of the SOAPy algorithm to predict INs.^61^ Given the dense arrangement of immune cells within the tonsil microenvironment and the average cell diameter of approximately 10 μm, we used a fixed radius r = 100 μm (including 10 cells within the diameter around the index cell) to ensure that spatial clustering analysis adequately covers cell neighbors’ signaling (including ligand-receptor binding and cytokine signaling). We conducted spatial clustering analysis on a tonsil spatial data that had undergone CmTSA multiplex staining and cell annotation (Fig. 5e), including 364,489 cells and 16 types of immune-related cells. All cells within the tissue were clustered into six INs (Fig. 8c). Then, with the assistance of pathologists, we manually analyzed the cellular components and spatial locations of each predicted INs and concluded that the six niches predicted by KNN spatial clustering were: IN-0) dark zone, IN-4) light zone, IN-2) interfollicular areas, IN-1) follicular mantle, IN-3) crypt, and IN-5) subepithelial regions (Fig. 8c).

Finally, paired cell-cell interaction analysis of the predominant cell types (comprising 90% of the cells in each niche) demonstrated that spatial interactions within the INs effectively reflect their immune functions (Fig. 8e). For example, in the crypt niche (IN-3), although epithelial cells were the main component, spatial interactions centered around DCs showed strong interactions with B cells, Th cells, and M1 macrophages, suggesting active antigen presentation and lymphocyte modulation in this region.

### Prediction and comparison of immune niches (INs) in patients at the whole-slide scale

Immune Niches (INs) reveal the immune activities occurring within the TME. Predicting INs at a whole-slide scale aids in dissecting the various immune activities and functions within patient’s TME, offering a potential strategy to inspect and quantify the heterogeneous immune status of individual patients.^17,62,63^ To test this method’s application, we conducted spatial clustering analysis on data derived from three cervical cancer patients who had undergone CmTSA multiplex staining and cell annotation (Cell number:CC_P1, 326,092; CC_P2, 295,578 ; CC_P3, 256,709) (Fig. 9a, Supplementary Fig. 7) . Our analysis revealed that, despite all patients having the same tumor type (cervical cancer), the frequencies of identified INs (eight in total) within their TMEs varied, indicating differences in their immune activities or status. Notably, we found that specific immune niches predominated in certain patients, potentially explaining the heterogeneity of immune activity among individual patients. For instance, IN-4 was predominantly present in CC_P1, mainly locating within the tumor stroma, closely surrounding and contacting the tumor mass. Unlike other immune niches, IN-4 was not dominated by a single cell type; its main components included dendritic cells (DCs), T helper cells (Th), plasma cells, B cells, cytotoxic T lymphocytes (CTLs), and PDL1+ tumor cells. Further analysis of cell-cell interactions within IN-4 suggested frequent interactions among DCs, Th, and B cells, indicating a dynamic immune response within the TME. However, significant interactions between PDL1+ tumor cells and CTLs also indicated ongoing immune suppression activities. In contrast, IN-7 was the specific and dominant INs in the TME of CC_P2. Spatially, IN-7 was located within the tumor stroma, closely surrounding the tumor mass and separating other lymphocyte-rich INs (such as IN-4 and IN-5) from contacting the tumor mass. The dominant cell type in IN-7 was FAP+ fibroblasts (Actived_fibro), known for their role in modulating ECM remodeling, secreting TGF-beta, and recruiting regulatory T cells (Tregs) and myeloid-derived suppressor cells (MDSCs).^64,65^ Thus, IN-7 may represent the main immune suppression mechanism in CC_P2. In CC_P3, both Niche-4 and Niche-7 were present in the TME, neither acting as the dominant niche, further indicating the heterogeneous immune activities occurring within individual TMEs. Overall, our results suggest that immune niche analysis can potentially identify and quantify various immune activities within the TME at a whole-slide scale. Certain IN can reveal the primary immune suppression activity happening in individual patients, highlighting the potential of this analysis for precision immunotherapy. However, these findings require careful validation in large clinical cohorts.

**Fig 9.**
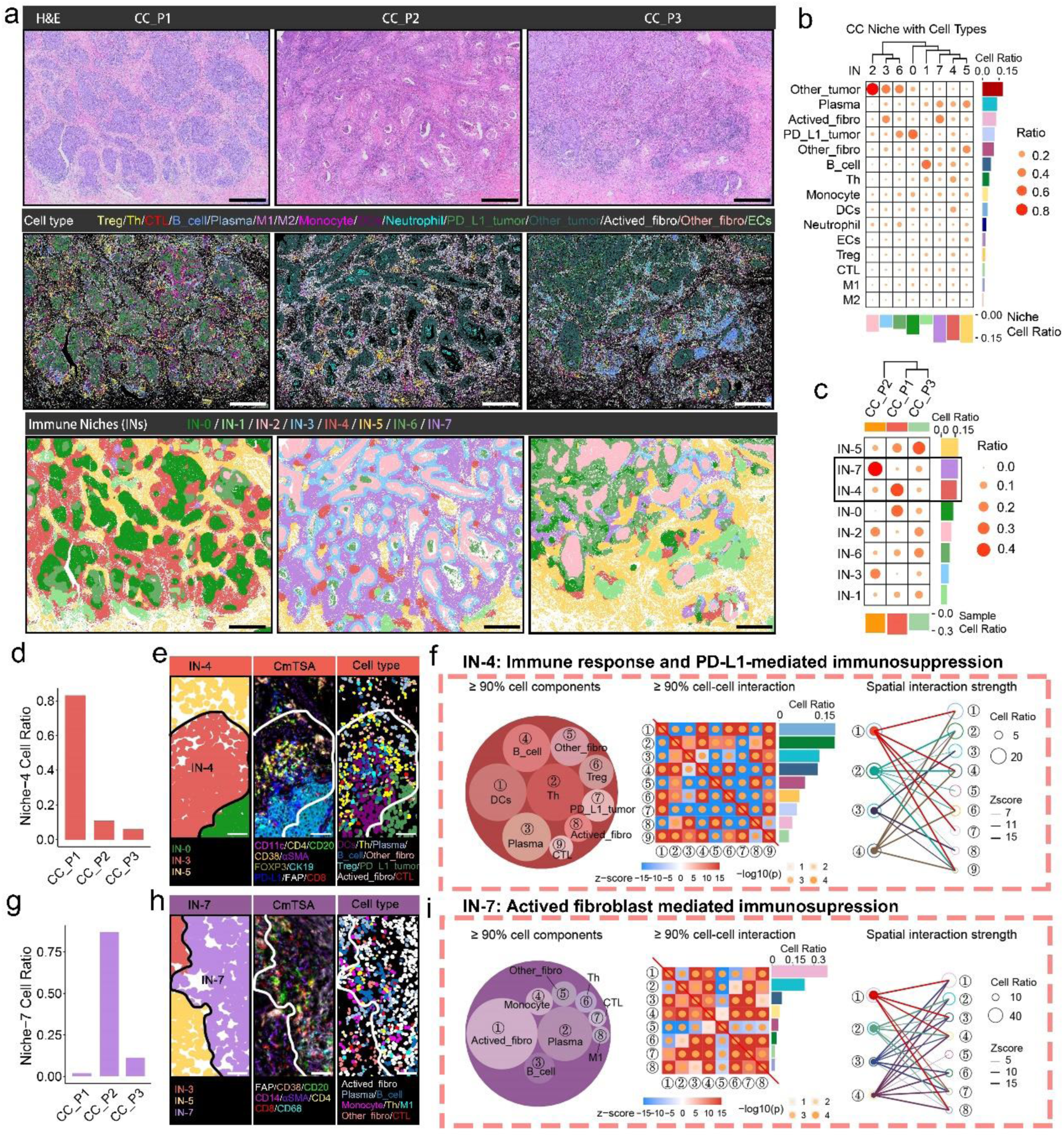
Prediction and comparison of immune niches in patients at the whole-slide scale. (a) TOP: Images of H&E staining from three cervical cancer patients; Middle: Cell type identification of the three cervical cancer patients based on cell type annotation rule; Bottom: Niche projection maps for the three cervical cancer patients. Scale bar, 1 mm. (b) Bubble plot of the cellular composition of the eight niches in the three cervical cancer samples, with color and size representing the ratio of cells in each niche. (c) Bubble plot of the niche composition in the three cervical cancer samples, with color and size representing the ratio of niches in each patient. (d) Bar chart of the ratio of IN-4 in the three patients. (e) Representative images: Left: IN -4 niche projection map; Middle: CmTSA staining image, showing only the top 90% of cells in IN-4; Bottom: Cell type map corresponding to the staining. Scale bar, 50 μm. (f) Left: Small circles represent the top 90% of cells in IN-4, with circle size indicating the relative proportion of each cell type; Middle: Heatmap of cell interactions, with the same meaning as in Fig. 7c, d; Right: Line plot showing interaction strength between the top 4 cell types by ratio and the remaining cells within the top 90%. Circle size represents cell ratio, and line thickness indicates interaction strength. (g) Bar chart of the ratio of IN-7 in the three patients. (h) Representative images: Left: IN-7 niche projection map; The remaining parts are identical to Fig. 9e. Scale bar, 50 μm. (i) Same as Fig. 9f, except the focus is on IN-7.

## DISCUSSION

Recent years, it has become increasingly appreciated that the features of individual TME are closely associated with tumor progression and response to treatment.^40^ TME mainly regarded to different cell types, their spatial distribution, cell-cell interaction and spatially structured arrangements (cellular niches or neighborhood).^39,41^ To capture the complexity of the TME, a suite of spatial profiling technologies have been developed, which can roughly divided into two categories: 1) spatial transcriptomics (10x Visium and Slide-seq) and 2) spatial proteomics (CODEX, MICS, MIBI-TOF and IMC).^4,11,14,66,67^ The application of these technologies has helped to unravel the structure and function of the tumor microenvironment, but there are several challenges need to overcome to unlock its full potential.

One of the greatest challenges of current spatial profiling technologies is the limited throughput for parallel sample testing.^15^ Cancer-related studies require to evaluate or compare cohort samples (generally containing several to dozens or even hundrens samples in each cohort size) to overcome inter-tumor and intra-tumor heterogeneity and obtain meaningful results.^68^ This is true for studies using animal models, and is dramatically exacerbated in studies with patients’ samples, where increased biological and treatment heterogeneity is compounded. The number of samples that can be uploaded during each experiment using current commercially available technologies is generally limited to just a few, typically 1-2 tissue slides per test.^26^ Here, we have developed the CmTSA method, an automated workflow that optimizes the superplex staining process. First, Abs and fluorophores are applied to samples using automated immunostaining equipment to ensure high-throughput labeling. Next, the stained tissue slides, up to dozens at a time, undergo imaging scanning to capture high-resolution images. Then, Abs are stripped, and fluorophores are quenched by our quenching system, preparing the samples for additional rounds of labeling.

Finally, serial images from iterative staining are registered into a cohesive dataset for analysis. This CmTSA workflow simplifies the experiment, increases batch size, and enhances consistency and processing speed, resulting in reliable and high-throughput outcomes for superplex immunostaining.

As previously mentioned, the complexity of the TME arises from the diversity of cell types, their spatial distribution, cell-cell interactions, and their structured spatial arrangements.^1,2,39^ Identifying cell identity and states within the TME using phenotype markers and co-expressed functional antigens is a fundamental step in spatial profiling analysis based on superplex immunostaining.

However, most microscopy-based superplex spatial proteomics methods (≥30 labeling) primarily use pre-labeled primary Abs with fluorophores or DNA oligos.^69^ The lack of a signal amplification step can result in dim fluorescence signals for cell markers, especially those with low expression levels, such as PD-1, CTLA-4, TIM-3, and FOXP3. TSA can amplify the signal of the primary Ab by catalyzing the deposition of tyramide-bound labels (such as fluorophores or enzymes) at the antigen site, which is particularly useful for detecting low-abundance targets.^13^ Here, we provide a quenching system capable of completely erasing TSA fluorescence signals within minutes. Thus, unlike other microscopy-based spatial proteomics platforms, CmTSA represents a superplex iterative staining method that is particularly suitable for detecting low-abundance antigens and rare cell types.

Another major limitation of superplex immunostaining is its high cost. This expense primarily stems from the instruments and reagents involved. To achieve full automation and lower technical barriers, manufacturers have integrated immunostaining, fluorophore signal recycling, and imaging scanning into a single machine. However, this integrated design strategy not only increases equipment costs but also reduces testing throughput due to hardware limitations.^11,13,70^ In contrast, CmTSA is an automated workflow that separates the superplex staining process into distinct steps: immunostaining, scanning, fluorophore quenching, and registration. Each step is completed using specific automated instruments, most of which are commonly available in pathology labs. This significantly reduces the cost of instruments. Moreover, because the CmTSA workflow is based on the TSA staining method, it is compatible with conventional IHC Abs and generally uses a lower concentration of primary Abs compared to immunostaining (IF) (either direct or indirect). Consequently, the CmTSA platform is a more cost-effective solution for superplex immunostaining.

Finally, and most importantly, superplex immunostaining enables the visualization of dozens of marker proteins across up to millions of cells within a whole-mount tissue slide. This vast pixel dataset presents multidimensional information that characterizes the patient’s TME. It includes multiple layers of information such as single-cell phenotypes, single-cell morphology, composition of cells in the tissue, cell-cell interactions, and their arrangements into multicellular niches or larger anatomical/functional structures.^68^ However, extracting these features from the raw pixel data is not straightforward, and requires dedicated analysis methods, bridging the worlds of microscopic images, pathology analysis and computer science. Here, following the CmTSA superplex immunostaining, we introduced an analysis pipeline to unravel the meaningful multidimensional spatial information embedded in the raw pixel data from the CmTSA staining.

This pipeline is logical and straightforward, starting with deep learning-based computer vision algorithms to segment individual cells from the large pixel-based image. Next, a rational cell type annotation rule is used to classify cells into different types. Based on the spatial matrix, the spatial distribution tendency of individual cell types, the interaction intensity of paired cells, and the multicellular functional niches are predicted and evaluated. By simply following this workflow, researchers can effectively visualize and quantify the types, states, and levels of immune activities within an individual patient’s immune microenvironment. This can significantly aid in basic research related to tumor immunology and in the analysis of immune microenvironment status for drug development and precision medicine.

While comprehensive spatial profiling of the TME is valuable and informative, it is essential to uncover the mechanisms behind the formation of spatial interactions and organizational patterns, as well as to reveal their functions and clinical implications. To address these questions, it is necessary to integrate spatial profiling technologies with multi-omics data, carefully annotated patient cohorts, and additional well-controlled clinical data. This integrated approach will provide deeper insights into the TME and its role in disease progression and treatment outcomes.

## MATERIALS AND METHODS

### Clinical samples

All clinical FFPE section samples, including those from tonsils, lung, colon, spleen, pancreas, stomach, kidney, liver, cervical cancer, and lung squamous cell carcinoma, were obtained from the Department of Pathology at Sichuan University West China Hospital. The use of these clinical samples was conducted in strict accordance with ethical guidelines and received approval from the Institutional Review Board (IRB) of Sichuan University West China Hospital.

This study involved four male patients, aged 55 to 57, all diagnosed with lung squamous cell carcinoma. Following their diagnosis, each patient received neoadjuvant chemotherapy. After the treatment, surgical resection specimens were obtained and pathologically examined, with all cases showing a pathological partial response (pPR). We performed dual immunofluorescence staining for CD8 and panCK on FFPE sections from these four patients. The study quantified the degree of aggregation of CTLs in different regions and the Infiltration Score of CTLs.

### Animal samples for mTSA

Adult C57 mice, housed in an SPF-grade animal facility, were anesthetized and perfused with 50 mL of PBS via the left ventricle, allowing blood to exit from the right atrium. Following this, the mice were perfused with 50 mL of 4% PFA (paraformaldehyde) tissue fixative. Various organs (brain, heart, liver and spleen) were then collected and immersed in 25 mL of 4% PFA for fixation. After 48 hours of fixation, the tissues were dehydrated and embedded. Tissue sections of 4 μm thickness were prepared using a paraffin microtome, floated on 45°C warm water to flatten, and then carefully transferred to the center of adhesive glass slides. The sections were baked at 60°C for one hour to ensure firm adherence to the slides.

### Cell line and mouse primary neuronal cells

HeLa cells were cultured in DMEM (Dulbecco’s Modified Eagle Medium) supplemented with 10% fetal bovine serum (FBS) and 1% penicillin-streptomycin. The cells were maintained in a humidified incubator at 37°C with 5% CO₂. For the preparation of coverslips, cells were seeded onto sterilized glass coverslips placed in 24-well culture plates and allowed to adhere for 24 hours prior to further experimental procedures.

For the mouse primary neuronal cells, Subventricular Zone (SVZ) neurons were extracted from postnatal day 8 (P8) mouse pups using a previously established protocol. Briefly, the brains were harvested and the SVZ was isolated, followed by enzymatic dissociation with acuutase and mechanical trituration to obtain a single-cell suspension. The cells were then cultured in Neurobasal medium supplemented with 2% B27, 0.5 mM L-glutamine, and 1% penicillin-streptomycin. The culture conditions were maintained at 37°C with 5% CO₂ in a humidified incubator. Similar to the HeLa cells, the primary neuronal cells were seeded onto sterilized glass coverslips in 24-well plates and allowed to adhere for 24 hours before proceeding with the subsequent experiments.

### Deparaffinization and antigen retrieval

Paraffin-embedded tissue sections were placed in a thermostatic incubator set at 65°C and incubated overnight. The sections were then sequentially immersed in staining jars containing xylene (three changes), with each immersion lasting 5 minutes. Following this, the sections were gradually rehydrated by immersing them in staining jars containing absolute ethanol (three changes), 95% ethanol, 85% ethanol, and 75% ethanol, each for 3 minutes. Finally, the sections were placed in a staining jar with purified water and rinsed three times, 3 minutes each. For antigen retrieval, the sections were placed in a staining jar containing 1x EDTA antigen retrieval solution and heated in a microwave until small bubbles formed in the solution. The jar was then transferred to a preheated water bath set at 95°C, where the sections were incubated for 30 minutes. Afterward, the sections were allowed to cool to room temperature naturally. The sections were then washed with PBS three times, 2 minutes each.

### H&E staining

After deparaffinization and antigen retrieval, the tissue sections are stained with hematoxylin, which binds to nucleic acids and colors the cell nuclei blue. Next, the sections undergo a bluing step, where they are treated with a weak alkaline solution to intensify the blue color of the hematoxylin. Following this, the sections are stained with eosin, which binds to cytoplasmic and extracellular proteins, providing a pink to red contrast. The sections are then dehydrated through a series of graded ethanol solutions, finishing in xylene. Finally, the stained sections are mounted with a coverslip using a permanent mounting medium. This process results in tissue sections with nuclei stained blue and cytoplasmic components stained in various shades of pink, allowing for detailed histological examination.

### Antibody infromations

See supplementary Table S1.

### Workflow of Cyclic mTSA

#### Background fluorescence quenching

After deparaffinization and antigen retrieval, the tissue sections were placed in the LUMINIRIS Fluorescence Quenching System (#MH030101). The sections were covered with an ample amount of IRISKit® HyperView quench buffer (#MH010301) and subjected to fluorescence quenching under 600W for 8 minutes to reduce background fluorescence. Following quenching, the sections were washed with PBS.

### Multiplex TSA (mTSA)

Before staining, a hydrophobic barrier was drawn around the tissue sections using an immunohistochemistry pen. The IRISKit® HyperView multiplex immunostaining kit (#MH010101) was used for antigen labeling. The sequential labeling was performed with primary antibody for 20 minutes, HRP-conjugated secondary antibody for 20 minutes, and fluorescent dye for 5-10 minutes per antibody. After each antibody labeling, the IRISKit® HyperView Advanced Ab Stripping Kit was used for antibody stripping. Different tyramide-based fluorescent dyes were used for each antibody staining, specifically Cyclic-480 (EX: 465-495; EM: 512-558), Cyclic-550 (EX: 540-560; EM: 575-595), and Cyclic-630 (EX: 620-640; EM: >665). After completing the three antibody labels, DAPI was used for nuclear staining. Between different reagents, the sections were washed twice with PBS, each wash lasting 3 minutes. Finally, the sections were mounted with a mounting medium and whole-slide imaging was performed using the EVIDENT VS200 microscope with a 20x objective lens.The image resolution was 0.345 μm/px.

### Differential quenching of fluorescent molecules

After imaging is completed, proceed with the quenching of the specific staining fluorescence, while ensuring that DAPI remains unquenched to be reserved as coordinates for image registration. The procedure and background fluorescence quenching are performed in the same manner. Once quenching is complete, proceed with the next round of staining using three antibodies.

### Image registration

After acquiring images from multiple rounds of imaging, an algorithm automatically identifies approximate feature points in the DAPI channel. A rigid transformation (translation, rotation) is then applied to find an optimal overlapping method that minimizes errors, ensuring the maximum overlap of all feature points. Subsequently, the images are standardized by converting the bit depth to a uniform 8-bit format with a grayscale range of 0 to 255. This standardization allows the images to be properly displayed on standard monitors, facilitating subsequent registration, analysis, and other operations.

### IF staining of FFPE sections

After completing antigen retrieval on paraffin sections, perform an 8-minute quenching of background fluorescence, followed by a 3-minute PBS wash, repeated twice. Incubate the sections with the appropriate primary antibody at 37°C for 1 hour. Afterward, wash with PBS for 3 minutes, repeated twice, and then add the corresponding fluorescent secondary antibody, incubating at 37°C for 30 minutes. Following this, wash with PBS for 3 minutes, repeated twice, and stain the nuclei with DAPI diluted 1:500 for 5 minutes. Wash again with PBS for 3 minutes, repeated twice, and mount the sections using a suitable mounting medium. Finally, perform whole-slide imaging using the EVIDENT VS200 microscope with a 20x objective lens.

### mTSA staining of cultured cells

Grow healthy HeLa cells or neuron-differentiated cells in a 12-well plate. Wash the cells with PBS for 3 minutes, repeating the wash twice. Fix the cells with 4% PFA at 4°C for 30 minutes. After fixation, wash the cells again with PBS for 3 minutes, repeating the wash twice. Add a sufficient amount of IRISKit® HyperView quench buffer (#MH010301) to cover the cells, and incubate for 5 minutes. Proceed with the antibody staining steps according to the mTSA protocol used for FFPE samples. Finally, perform imaging using the EVIDENT SpinSR microscope with a 100x objective lens.

### Cell segementation

The StarDist model is a powerful tool for accurately segmenting cell nuclei in microscopy images. We imported the pretrained StarDist model into QuPath for cell segmentation in multicolor images.

The preprocessed DAPI channel image was then passed through the StarDist model, which employs a star-convex polygon representation for precise nuclear segmentation. Simultaneously, we obtained the mean fluorescence intensity for each channel across different cellular structures, including nuclei, cytoplasm, membrane, and the whole cell, enabling us to filter marker positivity effectively. Additionally, we extracted the x and y coordinates of each cell’s nuclear centroid, which will be used to calculate spatial relationships between cells.

### Cell type Thresholds for marker positivity and lineage assignment

We identify the threshold for each marker channel individually in every multiplex image of FFPE sections. First, we select the channel for which the threshold needs to be set. Then, we choose the specific cellular location where the marker is expressed, such as the nucleus or membrane. A threshold is set within the 0-255 range, where cells with values equal to or above the threshold are classified as marker-positive, and those below the threshold are classified as marker-negative. This process is manually supervised, with marker-positive cells displayed on the stained image using a predefined mask color.Since certain antigen markers are shared by multiple cell types, and some cell types require multiple markers for accurate identification, an annotation rule is essential for ensuring consistent, accurate, and automated classification of cellular subtypes. To assign cellular phenotypes, we established a method based on typical lineage markers, expected population abundance, and staining quality supervision.

### Tissue region segmentation

For the tissue segmentation of cervical cancer stained images, tumor regions were outlined on the H&E images of adjacent sections under the guidance of a pathologist. These outlined regions were then loaded as annotations into the stained images in QuPath, allowing for the differentiation of tumor and stromal regions within the stained images.For patients with lung squamous cell carcinoma, Tumor regions were similarly outlined on the H&E images of adjacent sections under the pathologist’s guidance. These annotations were loaded into the stained images in QuPath, with the annotations extended outward by 100μm to define the Tumor-margin region, while the remaining area was designated as the Border region.

### Fluorescence intensity measurement

#### Background fluorescence quenching

Before and after quenching, use the same imaging parameters on the same section. Randomly select six non-overlapping ROIs (Regions of Interest) and measure the fluorescence intensity of three imaging channels using QuPath.

### Specific marker quenching efficiency

For five strictly consecutive sections, perform the same triple staining, using DAPI as the nuclear stain. After staining, perform whole-slide scanning under identical imaging conditions. The first section is not quenched, the second is quenched for 3 minutes, the third for 6 minutes, the fourth for 9 minutes, and the fifth for 12 minutes. After quenching, perform whole-slide scanning again under identical imaging conditions. At the same location on all five consecutive sections, randomly select six non-overlapping ROIs and measure the fluorescence intensity in the stained channels using QuPath.

### Antibody stripping

After performing the first antibody staining on the tissue (using the Cyclic-480 fluorophore) and completing the antibody stripping, the effectiveness of the stripping was tested by re-incubating the tissue with the secondary antibody for the first antigen for 20 minutes, followed by visualization with the Cyclic-550 fluorophore. After imaging, six non-overlapping ROIs were randomly selected in QuPath to measure the fluorescence intensity of Cyclic-480 and Cyclic-550.

### Signal-to-Noise Ratio of image

To calculate the Signal-to-Noise Ratio (SNR), we use the following formula:

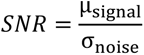

μ_signal_ is the mean pixel value of the signal region. σ_noise_ is the standard deviation of the noise region.

Use two strictly consecutive sections—one subjected to background fluorescence quenching and the other not. Perform antigen labeling under identical staining conditions. At the same location on both consecutive sections, randomly select six non-overlapping ROIs and measure the SNR in the stained channels using QuPath.

### Statistics of staining image

Statistical analyses were conducted using GraphPad Prism 10.1.2 software. Data are presented as mean ± standard error of the mean (SEM). To compare differences between two groups, an unpaired Student’s t-test was utilized. The p-values were interpreted as follows:ns: p > 0.05 (not statistically significant); *: p ≤ 0.05; **: p ≤ 0.01; ***: p ≤ 0.001; ****: p ≤ 0.0001. These symbols indicate the level of statistical significance, with more asterisks representing a higher level of significance.

### Spatial aggregation

For each cell within the region of interest in an image, we calculated the average distance to the five nearest neighboring cells of the same type. This average distance was then divided by the sparsest distance of the same cell type. The sparsest distance of the same cell type was calculated by dividing the total number of cells of that type by the total area of the region. This standardized reference value allows for a more comparable measurement of cell spatial aggregation across different regions or samples, providing a clearer expression of the observed aggregation.

After implementing the calculation of spatial aggregation in the R (4.3.1) environment, we normalized the data within the same patient to a range between 0 and 1 (using the formula below). First, we applied the arctangent function (*atan*) to each aggregation value for nonlinear transformation. This operation smooths the data, reducing the impact of noise or irregularities on the results, especially when dealing with noisy data, providing more stable analysis outcomes. Then, we calculated the minimum value min(atan(*A*)) and the maximum value max(atan(*A*))of the transformed *atan* data, representing the minimum and maximum range of the data, respectively. Finally, the atan-transformed data was normalized by linearly mapping it to the range between 0 and 1, where values closer to 1 indicate weaker spatial aggregation of that cell type, and values closer to 0 indicate stronger aggregation.

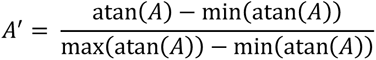

In the formula above, *A* represents the original aggregation value, *atan* denotes the arctangent function, and *A^′^* represents the normalized aggregation value.

In Fig. 6b, we compared the aggregation of each cell type between the normal group and the permutation group. The data for the permutation group was generated by randomizing the coordinates in the real data and then extracting the current cell type as the permutation data. This process was repeated 10 times, and the mean was taken as the aggregation value for each cell in the permutation group, which was then compared with the normal group.

### Infiltration Score

To assess the infiltration of CTLs within the tumor region, we introduced an Infiltration Score (IS). This score is calculated by connecting all cells within a 50μm radius of the tumor cells and then determining the number of connections between different types of cells. The Infiltration Score is computed using the following formula:

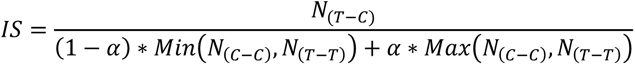

Where *N*_(*T*−*C*)_ represents the number of connections between tumor cells and CTLs, *N*_(*C*−*C*)_ represents the number of connections between CTLs, and *N*_(*T*−*T*)_ represents the number of connections between tumor cells themselves. *Min* represents the minimum value between *N*_(*C*−*C*)_ and *N*_(*T*−*T*)_, *Max* represents the maximum value between *N*_(*C*−*C*)_ and *N*_(*T*−*T*)_. The parameter α is a weighting factor used to reduce the impact of extreme values while preserving the influence of larger values, thereby stabilizing the data results. In this study, focused on lung squamous cell carcinoma samples, we set the value of α to 0.1.

### Construction of the spatial network

Before conducting spatial analysis, we needed to construct a spatial network of cells. For the initial visualization of spatial relationships between cells, we first used the nn2 function from the RANN (2.6.1) package in the R (4.3.1) environment to construct a preliminary spatial network. We set the parameters as *k* = 15 and searchtype = "*radius*", and defined a radius of 15 μm (*radius* = 15) to determine neighborhood relationships. In this process, the focal cell itself was excluded when selecting neighbors, considering only the first 15 neighboring cells within the 15 μm radius to establish network connections. The resulting network graph provided an initial visualization of the spatial distribution of cells and their adjacency relationships, offering an intuitive preliminary view for further analysis.

For more detailed spatial analysis, we employed the Spatial Network Construction method from the SOAPy package. This approach allowed us to analyze interactions between cell types at a finer scale, thereby revealing more complex biological phenomena.

### Spatial interaction analysis

To identify significant interactions and separations between cell types, we used the Spatial Neighborhood Analysis method from SOAPy. All calculations and plotting were completed in Python (3.9) and R (4.3.1). The core of this method involves calculating the number of edges between each pair of cell types in the network and comparing it with the sum of edges between the paired cell types and other cells. During the calculation, a permutation test was introduced to assess the specificity of the distribution in the real data. Specifically, this involved generating *n* = 1000 randomizations of cell distributions and calculating the interaction *Z-score* between each pair of cell types. By converting the *Z-scores* into *P-values*, we could determine whether the interactions between cell types were statistically significant. This process provides a reliable statistical basis for the biological interpretation of cell type interactions and subsequent research. The calculation process is as follows:

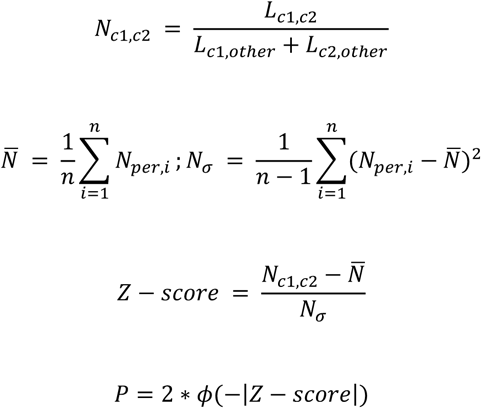

In the formula above, *L*_*c*1,*c*2_ represents the number of edges between cell types *c*1 and *c*2; *L*_*c*1,other_ represents the number of edges between *c*1 and all other cell types; *L*_*c*2,other_ represents the number of edges between *c*2 and other cell types; *N*_*c*1,*c*2_ represents the cell interaction score; *N_per,i_* represents the number of edges for the cell type pair in the *i* th permutation test; *N* represents the average number of edges for the specific cell type pair across all permutation tests; and *N*_α_ represents the overall standard deviation, which quantifies the dispersion of the number of edges for the specific cell type pair in the permutation tests. The *Z-score* is a standardized measure indicating the deviation of the observed value from the mean, where a positive *Z-score* suggests a tendency for interaction between cell types, and a negative *Z-score* suggests a tendency for separation. In the heatmap visualization, *Z-scores* are limited to the range of -15 to 15 to better display the results. *Z-scores* exceeding this range are truncated to 15 or -15, respectively, to avoid extreme values affecting the chart, enhancing readability and analytical validity.

Additionally, we used the *P-values* converted from the *Z-scores* in the heatmap to provide statistical confidence for deviations. By calculating *-log10(P)*, *P-values* are transformed into a more intuitive form and overlaid on the heatmap. This transformation allows the heatmap to not only display the magnitude and direction of *Z-scores* but also reflect the degree of statistical significance, providing a clearer understanding of the reliability and importance of the results.

### Spatial functional domain analysis

We used the Spatial Composition Analysis method from SOAPy. In this method, we first define the ratio of surrounding cells for each index cell, and groups of index cells with similar surrounding cell compositions are defined as a niche (see formula below). After identifying each niche using *K-means* clustering, we visualized the data in R (4.3.1).

Specifically, we used ggplot (3.4.3) to visualize the projection maps of cell types and niches. Next, we computed the Euclidean distance matrix using the dist function from the Stat (4.3.1) package and performed hierarchical clustering using the hclust function from the flashClust (1.01-2) package, generating a dendrogram. Finally, we displayed the cell composition in each niche using a bubble plot, allowing us to clearly observe the cell components and characteristics of different niches.

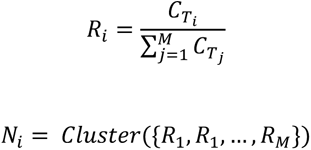

In the formula, 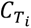 represents different cell types for the index cell ; 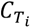 represents the number of cells of the surrounding cell; *M* represents the total number of cell types; 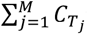 represents the total number of surrounding cells for the index cell; and *R_i_* indicates the ratio of cells of type *T_i_* in the niche of the index cell.

### Statistics and plotting of spatial analysis

All statistical plots were generated using ggplot2 (3.4.3) in the R (4.3.1) environment. Data were preprocessed and cleaned before plotting. Any statistical analyses and tests were completed prior to plotting. Custom themes and settings were applied to the charts. Specific codes and scripts are available to ensure the reproducibility of the results.

## DATA AVAILABILITY

The original data can be obtained by contacting the corresponding author, Chengjian Zhao. Additional data can be found in the MATERIALS AND METHODS section. All computer codes used for analyzing spatial distributions, along with further details, are available from the corresponding author upon request.

## ACKNOWLEDGEMENTS

This research was supported by various funding sources, including the National Natural Science Foundation of China (grant no. 8227054), Natural Science Foundation of Sichuan Province (grant no. 2023NSFSC0665, 2023NSFSC1833, 2023NSFSC0539), the National Key Scientific and Technological Project of Ningxia Medical University (grant no. 23H1529), and the National Key Research and Development Program of China (grant no. 2023YFB330850103).

## AUTHOR CONTRIBUTIONS

Chaoxin Xiao, Ruihan Zhou, and Qin Chen contributed equally to this work. Chaoxin Xiao and Qin Chen were responsible for the conception and design of the study. Ruihan Zhou and Qin Chen performed the experiments. Wanting Hou, Xiaoying Li and Yulin Wang carried out the data analysis. Lu Liu, Huanhuan Wang, and Xiaohong Yao contributed to the acquisition of data.

Tongtong Xu and Fujun Cao provided critical revisions and helped with data interpretation. Banglei Yin and Ouying Yan contributed to the drafting of the manuscript. Lili Jiang and Wei Wang provided technical support and supervision. Chaoxin Xiao was also responsible for figure preparation. Dan Cao supervised the project and provided critical guidance throughout the study. Chengjian Zhao oversaw the entire project and was responsible for the final approval of the manuscript.

## REFERENCES

1. Arneth, B. Tumor Microenvironment. Medicina (Kaunas). 56, (2019).

2. Zhang, X., et al. The Tumor Microenvironment: Signal Transduction. Biomolecules. 14, (2024).

3. Pham, D., et al. Robust mapping of spatiotemporal trajectories and cell-cell interactions in healthy and diseased tissues. Nature communications. 14, 7739 (2023).

4. Goltsev, Y., et al. Deep Profiling of Mouse Splenic Architecture with CODEX Multiplexed Imaging. Cell. 174, 968–981.e915 (2018).

5. Fu, T., et al. Spatial architecture of the immune microenvironment orchestrates tumor immunity and therapeutic response. J Hematol Oncol. 14, 98 (2021).

6. Lei, X., et al. Immune cells within the tumor microenvironment: Biological functions and roles in cancer immunotherapy. Cancer letters. 470, 126–133 (2020).

7. Wang, N., Li, X., Wang, R., and Ding, Z. Spatial transcriptomics and proteomics technologies for deconvoluting the tumor microenvironment. Biotechnol J. 16, e2100041 (2021).

8. Xie, F., et al. Progress in research on tumor microenvironment-based spatial omics technologies. Oncol Res. 31, 877–885 (2023).

9. Denisenko, E., et al. Spatial transcriptomics reveals discrete tumour microenvironments and autocrine loops within ovarian cancer subclones. Nature communications. 15, 2860 (2024).

10. Anderson, A.C., et al. Spatial transcriptomics. Cancer cell. 40, 895–900 (2022).

11. Giesen, C., et al. Highly multiplexed imaging of tumor tissues with subcellular resolution by mass cytometry. Nature methods. 11, 417–422 (2014).

12. Lin, J.R., et al. Highly multiplexed immunofluorescence imaging of human tissues and tumors using t-CyCIF and conventional optical microscopes. eLife. 7, (2018).

13. Hoyt, C.C. Multiplex Immunofluorescence and Multispectral Imaging: Forming the Basis of a Clinical Test Platform for Immuno-Oncology. Frontiers in molecular biosciences. 8, 674747 (2021).

14. Angelo, M., et al. Multiplexed ion beam imaging of human breast tumors. Nature medicine. 20, 436–442 (2014).

15. Mund, A., Brunner, A.D., and Mann, M. Unbiased spatial proteomics with single-cell resolution in tissues. Mol Cell. 82, 2335–2349 (2022).

16. Chen, H.Y., Palendira, U., and Feng, C.G. Navigating the cellular landscape in tissue: Recent advances in defining the pathogenesis of human disease. Comput Struct Biotechnol J. 20, 5256–5263 (2022).

17. Schürch, C.M., et al. Coordinated Cellular Neighborhoods Orchestrate Antitumoral Immunity at the Colorectal Cancer Invasive Front. Cell. 182, 1341–1359.e1319 (2020).

18. Stringer, C., Wang, T., Michaelos, M., and Pachitariu, M. Cellpose: a generalist algorithm for cellular segmentation. Nature methods. 18, 100–106 (2021).

19. Schmidt, U., Weigert, M., Broaddus, C., and Myers, G. (2018). Cell detection with star-convex polygons. (Springer), pp. 265–273.

20. Stevens, M., et al. StarDist Image Segmentation Improves Circulating Tumor Cell Detection. Cancers. 14, (2022).

21. Schapiro, D., et al. histoCAT: analysis of cell phenotypes and interactions in multiplex image cytometry data. Nature methods. 14, 873–876 (2017).

22. Dries, R., et al. Giotto: a toolbox for integrative analysis and visualization of spatial expression data. Genome biology. 22, 78 (2021).

23. Lee, R.Y., et al. The promise and challenge of spatial omics in dissecting tumour microenvironment and the role of AI. Front Oncol. 13, 1172314 (2023).

24. Monici, M. Cell and tissue autofluorescence research and diagnostic applications. Biotechnology annual review. 11, 227–256 (2005).

25. Lin, J.R., Fallahi-Sichani, M., Chen, J.Y., and Sorger, P.K. Cyclic Immunofluorescence (CycIF), A Highly Multiplexed Method for Single-cell Imaging. Current protocols in chemical biology. 8, 251–264 (2016).

26. Willemsen, M., et al. Improvement of Opal Multiplex Immunofluorescence Workflow for Human Tissue Sections. The journal of histochemistry and cytochemistry : official journal of the Histochemistry Society. 69, 339–346 (2021).

27. Xu, N., et al. Downregulation of N4-acetylcytidine modification in myeloid cells attenuates immunotherapy and exacerbates hepatocellular carcinoma progression. British journal of cancer. 130, 201–212 (2024).

28. Tsurui, H., et al. Seven-color fluorescence imaging of tissue samples based on Fourier spectroscopy and singular value decomposition. The journal of histochemistry and cytochemistry : official journal of the Histochemistry Society. 48, 653–662 (2000).

29. Seo, J.H., et al. Automated stitching of microscope images of fluorescence in cells with minimal overlap. Micron (Oxford, England : 1993). 126, 102718 (2019).

30. Wodzinski, M., Marini, N., Atzori, M., and Müller, H. RegWSI: Whole slide image registration using combined deep feature- and intensity-based methods: Winner of the ACROBAT 2023 challenge. Computer methods and programs in biomedicine. 250, 108187 (2024).

31. Haghofer, A., et al. Histological classification of canine and feline lymphoma using a modular approach based on deep learning and advanced image processing. Scientific reports. 13, 19436 (2023).

32. Ourselin, S., et al. The 19th International Conference on Medical Image Computing and Computer-Assisted Intervention (MICCAI 2016). Medical image analysis. 41, 1 (2017).

33. Bankhead, P., et al. QuPath: Open source software for digital pathology image analysis. Scientific reports. 7, 16878 (2017).

34. Low, V., Li, Z., and Blenis, J. Metabolite activation of tumorigenic signaling pathways in the tumor microenvironment. Science signaling. 15, eabj4220 (2022).

35. Shan, F., et al. Therapeutic targeting of regulatory T cells in cancer. Trends in cancer. 8, 944–961 (2022).

36. van Vlerken-Ysla, L., Tyurina, Y.Y., Kagan, V.E., and Gabrilovich, D.I. Functional states of myeloid cells in cancer. Cancer cell. 41, 490–504 (2023).

37. Huang, J., et al. Extracellular matrix and its therapeutic potential for cancer treatment. Signal transduction and targeted therapy. 6, 153 (2021).

38. Xia, L., et al. The cancer metabolic reprogramming and immune response. Molecular cancer. 20, 28 (2021).

39. Pitt, J.M., et al. Targeting the tumor microenvironment: removing obstruction to anticancer immune responses and immunotherapy. Annals of oncology : official journal of the European Society for Medical Oncology. 27, 1482–1492 (2016).

40. Hanahan, D., and Coussens, L.M. Accessories to the crime: functions of cells recruited to the tumor microenvironment. Cancer cell. 21, 309–322 (2012).

41. Quail, D.F., and Joyce, J.A. Microenvironmental regulation of tumor progression and metastasis. Nature medicine. 19, 1423–1437 (2013).

42. Posselt, R., et al. Spatial distribution of FoxP3+ and CD8+ tumour infiltrating T cells reflects their functional activity. Oncotarget. 7, 60383–60394 (2016).

43. Yin, J., et al. Neoadjuvant adebrelimab in locally advanced resectable esophageal squamous cell carcinoma: a phase 1b trial. Nature medicine. 29, 2068–2078 (2023).

44. Tumeh, P.C., et al. PD-1 blockade induces responses by inhibiting adaptive immune resistance. Nature. 515, 568–571 (2014).

45. Fu, C., and Jiang, A. Dendritic Cells and CD8 T Cell Immunity in Tumor Microenvironment. Frontiers in immunology. 9, 3059 (2018).

46. Blanco, M., et al. Unveiling the Role of the Tumor Microenvironment in the Treatment of Follicular Lymphoma. Cancers. 14, (2022).

47. Tanaka, A., and Sakaguchi, S. Targeting Treg cells in cancer immunotherapy. European journal of immunology. 49, 1140–1146 (2019).

48. Shimasaki, N., Jain, A., and Campana, D. NK cells for cancer immunotherapy. Nature reviews. Drug discovery. 19, 200–218 (2020).

49. Vitale, I., et al. Macrophages and Metabolism in the Tumor Microenvironment. Cell metabolism. 30, 36–50 (2019).

50. Janesick, A., et al. High resolution mapping of the tumor microenvironment using integrated single-cell, spatial and in situ analysis. Nature communications. 14, 8353 (2023).

51. Hickey, J.W., et al. T cell-mediated curation and restructuring of tumor tissue coordinates an effective immune response. Cell reports. 42, 113494 (2023).

52. Grünwald, B.T., et al. Spatially confined sub-tumor microenvironments in pancreatic cancer. Cell. 184, 5577–5592.e5518 (2021).

53. Toda, S., Frankel, N.W., and Lim, W.A. Engineering cell-cell communication networks: programming multicellular behaviors. Current opinion in chemical biology. 52, 31–38 (2019).

54. Kamali, A.N., Bautista, J.M., Eisenhut, M., and Hamedifar, H. Immune checkpoints and cancer immunotherapies: insights into newly potential receptors and ligands. Therapeutic advances in vaccines and immunotherapy. 11, 25151355231192043 (2023).

55. Wang, J., et al. Unleashing the power of immune checkpoints: Post-translational modification of novel molecules and clinical applications. Cancer letters. 588, 216758 (2024).

56. Leonard, W.J., and Lin, J.X. Strategies to therapeutically modulate cytokine action. Nature reviews. Drug discovery. 22, 827–854 (2023).

57. Thurley, K., Gerecht, D., Friedmann, E., and Höfer, T. Three-Dimensional Gradients of Cytokine Signaling between T Cells. PLoS computational biology. 11, e1004206 (2015).

58. Mi, H., et al. Spatial and Compositional Biomarkers in Tumor Microenvironment Predicts Clinical Outcomes in Triple-Negative Breast Cancer. bioRxiv : the preprint server for biology. (2023).

59. García-Ortiz, A., et al. The Role of Tumor Microenvironment in Multiple Myeloma Development and Progression. Cancers. 13, (2021).

60. Nabhan, M., et al. Deciphering the tumour immune microenvironment cell by cell. Immuno-oncology technology. 18, 100383 (2023).

61. Wang, H., et al. SOAPy: a Python package to dissect spatial architecture, dynamics and communication. 2023.2012. 2021.572725 (2023).

62. Sorin, M., et al. Single-cell spatial landscapes of the lung tumour immune microenvironment. Nature. 614, 548–554 (2023).

63. Karimi, E., et al. Single-cell spatial immune landscapes of primary and metastatic brain tumours. Nature. 614, 555–563 (2023).

64. Croizer, H., et al. Deciphering the spatial landscape and plasticity of immunosuppressive fibroblasts in breast cancer. Nature communications. 15, 2806 (2024).

65. Qin, P., et al. Cancer-associated fibroblasts undergoing neoadjuvant chemotherapy suppress rectal cancer revealed by single-cell and spatial transcriptomics. Cell reports. Medicine. 4, 101231 (2023).

66. Du, M.R., et al. Spotlight on 10x Visium: a multi-sample protocol comparison of spatial technologies. 2024.2003. 2013.584910 (2024).

67. Rodriques, S.G., et al. Slide-seq: A scalable technology for measuring genome-wide expression at high spatial resolution. 363, 1463–1467 (2019).

68. Elhanani, O., Ben-Uri, R., and Keren, L. Spatial profiling technologies illuminate the tumor microenvironment. Cancer cell. 41, 404–420 (2023).

69. Lewis, S.M., et al. Spatial omics and multiplexed imaging to explore cancer biology. Nature methods. 18, 997–1012 (2021).

70. Balzarotti, F., et al. Nanometer resolution imaging and tracking of fluorescent molecules with minimal photon fluxes. Science. 355, 606–612 (2017).

