## Supplemental Table and Figures for "Pipeline for Assessing Tumor Immune Status Using Superplex Immunostaining and Spatial Immune Interaction Analysis": Supplementary-20240823.pdf

Supplementary Table 1. Antibody informations

| Antibody | Brand | Catalog# | Source | Clone number | Dilution rate |
|---|---|---|---|---|---|
| CD3 | HUABIO | HA720082 | Rabbit | JE80-02 | 1: 1000 |
| aSMA | HUABIO | ET1607-53 | Rabbit | SY02-64 | 1 : 3000 |
| Bax | CST | 5023 | Rabbit | D2E11 | 1: 1000 |
| BCL2 | abcam | ab241548 | Rat | NOR 235J | 1: 800 |
| BCL6 | HUABIO | HA601083 | Rabbit | A6C7-R | 1: 300 |
| CD11b | HUABIO | EM1701-42 | Mouse | 4-6 | 1: 500 |
| CD11c | HUABIO | ET1606-19 | Rabbit | SI19-06 | 1: 1000 |
| CD14 | HUABIO | ET1610-85 | Rabbit | SC69-02 | 1: 1000 |
| CD16 | abcam | ab308607 | Rat | SP175 | 1: 500 |
| CD163 | abcam | ab289979 | Rat | EPR19518 | 1: 2000 |
| CD163 | HUABIO | ET1704-43 | Rabbit | JA51-30 | 1: 2000 |
| CD20 | abcam | ab279300 | Rat | EP459Y | 1: 1000 |
| CD20 | HUABIO | HA721138 | Rabbit | PD00-02 | 1: 5000 |
| CD21 | HUABIO | HA721163 | Rabbit | PD00-23 | 1: 1000 |
| CD23 | HUABIO | HA721139 | Rabbit | PD00-03 | 1: 1000 |
| CD3 | abcam | ab11089 | Rat | CD3-12 | 1: 1000 |
| CD31 | HUABIO | M1511-8 | Mouse | 7-A1 | 1: 2000 |
| CD31 | abcam | ab182981 | Rabbit | EPR17259 | 1: 2000 |
| CD34 | HUABIO | ET1606-11 | Rabbit | SI16-01 | 1: 1000 |
| CD38 | HUABIO | HA721268 | Rabbit | PD01-49 | 1: 1500 |
| CD4 | HUABIO | ET1609-52 | Rabbit | ST0488 | 1: 1500 |
| CD57 | HUABIO | HA601114 | Rabbit | PD00-21 | 1: 1000 |
| CD68 | HUABIO | HA601115 | Mouse | PDM0-13 | 1: 2000 |
| CD8 | Immunoway | ABT304 | Mouse | ABT304 | 1: 500 |
| CD8 | Servicebio | GB114196 | Rabbit | Poly | 1: 500 |
| CK19 | HUABIO | ET1601-6 | Rabbit | SA30-06 | 1: 1000 |
| Ecad | HUABIO | HA601143 | Mouse | A0-G11-2-R | 1: 4000 |
| Ecad | HUABIO | ET1607-75 | Rabbit | SY0287 | 1: 1000 |
| EGFR | HUABIO | ET1604-44 | Rabbit | SP00-86 | 1: 1000 |
| F4/80 | Servicebio | GB113373 | Rabbit | Poly | 1: 500 |
| FAK | HUABIO | ET1602-25 | Rabbit | SR46-04 | 1: 1000 |
| FAP | zenbio | R381838 | Rabbit | R08-9H6 | 1: 1000 |
| FOXP3 | abcam | ab20034 | Mouse | 236A/E7 | 1: 800 |
| GrB | abcam | ab289888 | Rat | EPR8260 | 1: 100 |
| IDO1 | HUABIO | HA721331 | Rabbit | PD00-62 | 1: 800 |
| Ki67 | HUABIO | HA721115 | Rabbit | SR00-02 | 1: 3000 |
| laminA/C | HUABIO | ET7110-12 | Rabbit | JE51-60 | 1: 500 |
| MAP2 | HUABIO | HA500177 | Rabbit | Poly | 1: 500 |
| MPO | abcam | ab300650 | Rat | EPR20257 | 1: 4000 |
| MPO | HUABIO | ab208670 | Rabbit | EPR20257 | 1: 4000 |
| NeuN | HUABIO | ET1602-12 | Rabbit | SR45-07 | 1: 500 |
| panCK | HUABIO | HA601094 | Mouse | PD00-15 | 1: 500 |

| Antibody | Brand | Catalog# | Source | Clone number | Dilution rate |
|---|---|---|---|---|---|
| panCK | HUABIO | HA601138 | Mouse | PDH09-10 | 1: 3000 |
| PD1 | ZSGB-BIO | ZM-0381 | Mouse | UMAB199 | 1: 800 |
| PD-L1 | abcam | ab279294 | Rabbit | CAL10 | 1: 1000 |
| PS6 | CST | 4858 | Rabbit | D57.2.2E | 1: 1000 |
| S100A9 | HUABIO | ET1702-73 | Rabbit | JF096-8 | 1: 3000 |
| Tomm20 | HUABIO | ET1609-25 | Rabbit | ST04-72 | 1: 2000 |
| Tryptase | HUABIO | ET1610-64 | Rabbit | SC68-07 | 1: 3000 |
| Tuj1 | HUABIO | M0805-8 | Mouse | A8-D10 | 1: 500 |
| YAP | CST | 8418 | Rabbit | D24E4 | 1: 1000 |
| ZO1 | Proteintech | 21773-1-AP | Rabbit | Poly | 1: 1000 |

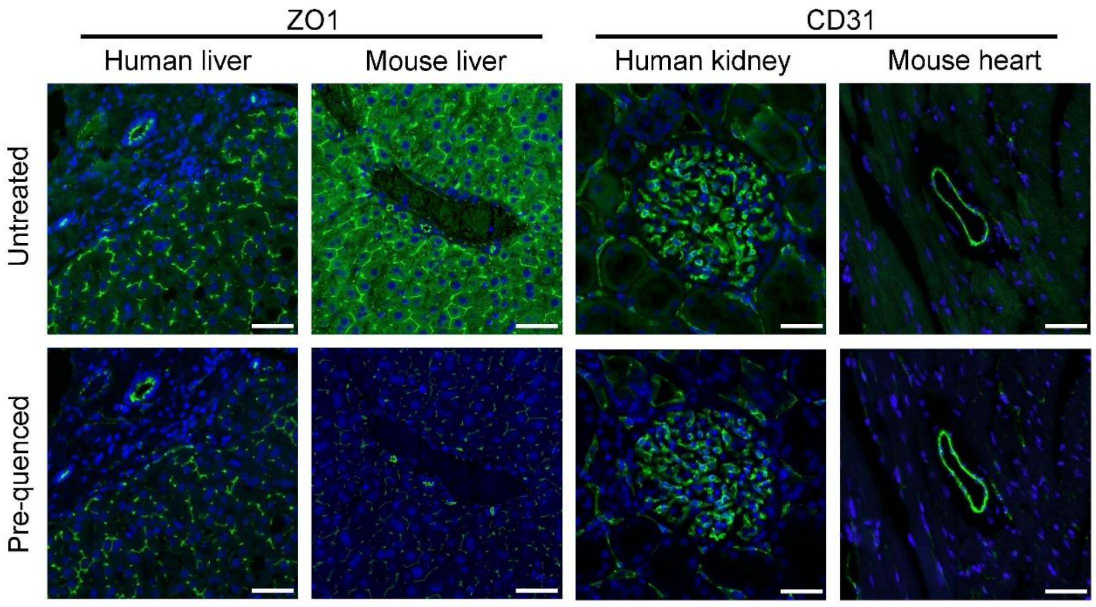


**Supplementary Fig. 1 Quenching background fluorescence could dramatically enhance the staining SNR.**

Representative images of different antibody markers after background fluorescence quenching. Scale bar, 50 μm.

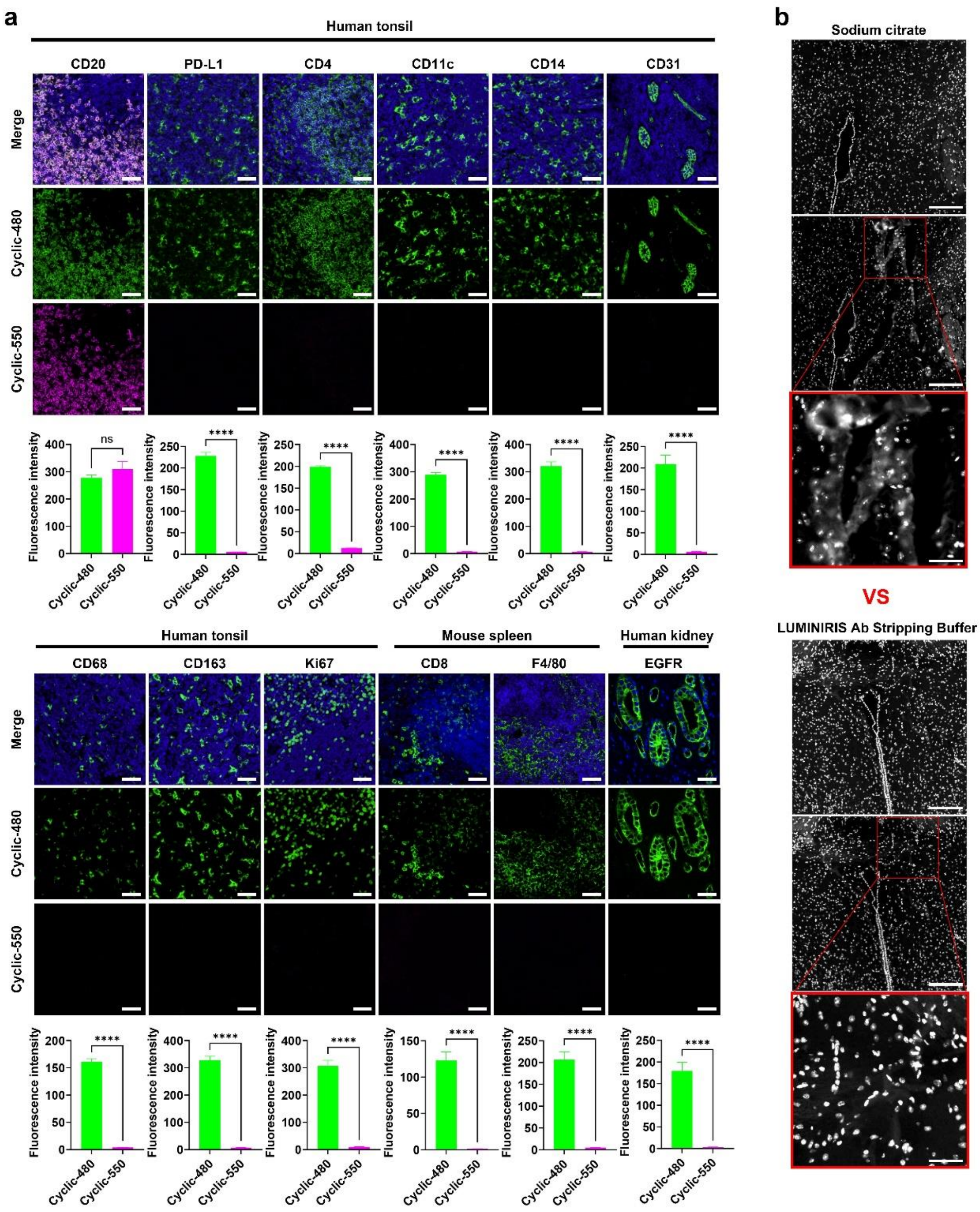


**Supplementary Fig. 2 LUMINIRIS Ab Stripping Buffer enables efficient antibody stripping without damaging tissue.**

(a) Efficient antibody stripping was achieved across various tissues for multiple antibodies. Specifically, CD20 was not stripped, and the fluorescence intensity of Cyclic-550 matched that of Cyclic-480. All other antibodies were efficiently stripped, with the fluorescence intensity of Cyclic-550 approaching zero. Scale bar, 50 μm.

(b) Mouse brain sections were subjected to two rounds of stripping using either sodium citrate retrieval solution or LUMINIRIS Ab Stripping Buffer, followed by nuclear staining with DAPI. The results indicate that the traditional sodium citrate retrieval method causes significant tissue damage, whereas LUMINIRIS Ab Stripping Buffer provides a gentler stripping, preserving the structural integrity of the tissue. Scale bar, 100 μm.

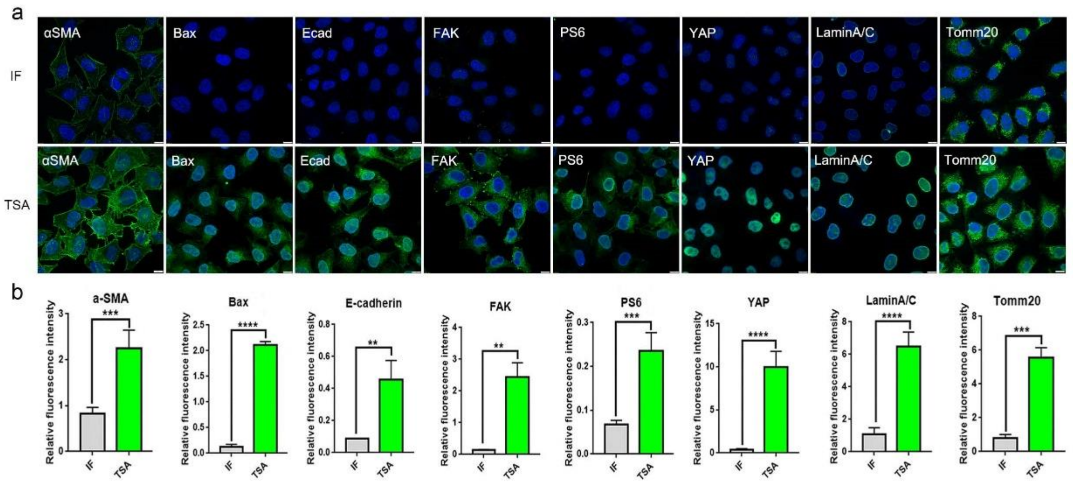


**Supplementary Fig. 3 The fluorescence intensity of the TSA method is 5 to 100 times higher than that of traditional IF staining.**

(a) Representative images of different types of antigens (cytoskeleton, transcription factors, membrane proteins, kinases, etc.) stained on Hela cells. Scale bar, 10 μm.

(b) Comparative statistical analysis of fluorescence intensity values between IF staining and TSA staining. Data are shown as Mean ± SEM. based on Unpaired Student's t-test. **: p ≤ 0.01; ***: p ≤ 0.001; ****: p ≤ 0.0001.

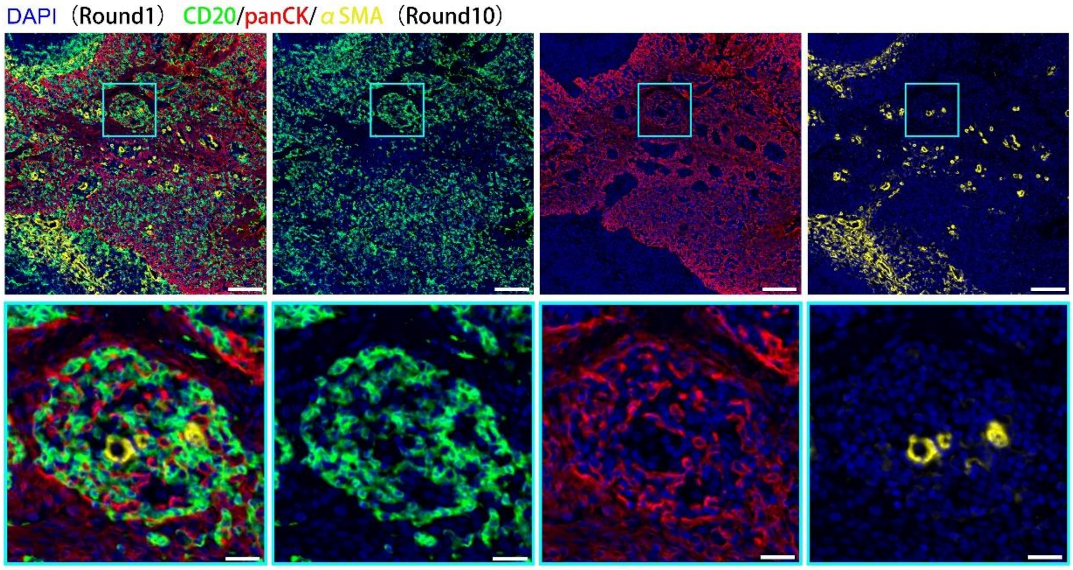


**Supplementary Fig. 4 Perfect pixel-to-pixel matching of fluorophore signaling to the first DAPI.**

Image showing the first-round DAPI and the 10th-round marker from the 30-marker panel in Fig. 4c. The results demonstrate a perfect match between the 10th-round marker and the first-round DAPI. Scale bar, 200 μm. Zoom in scale bar, 40 μm.

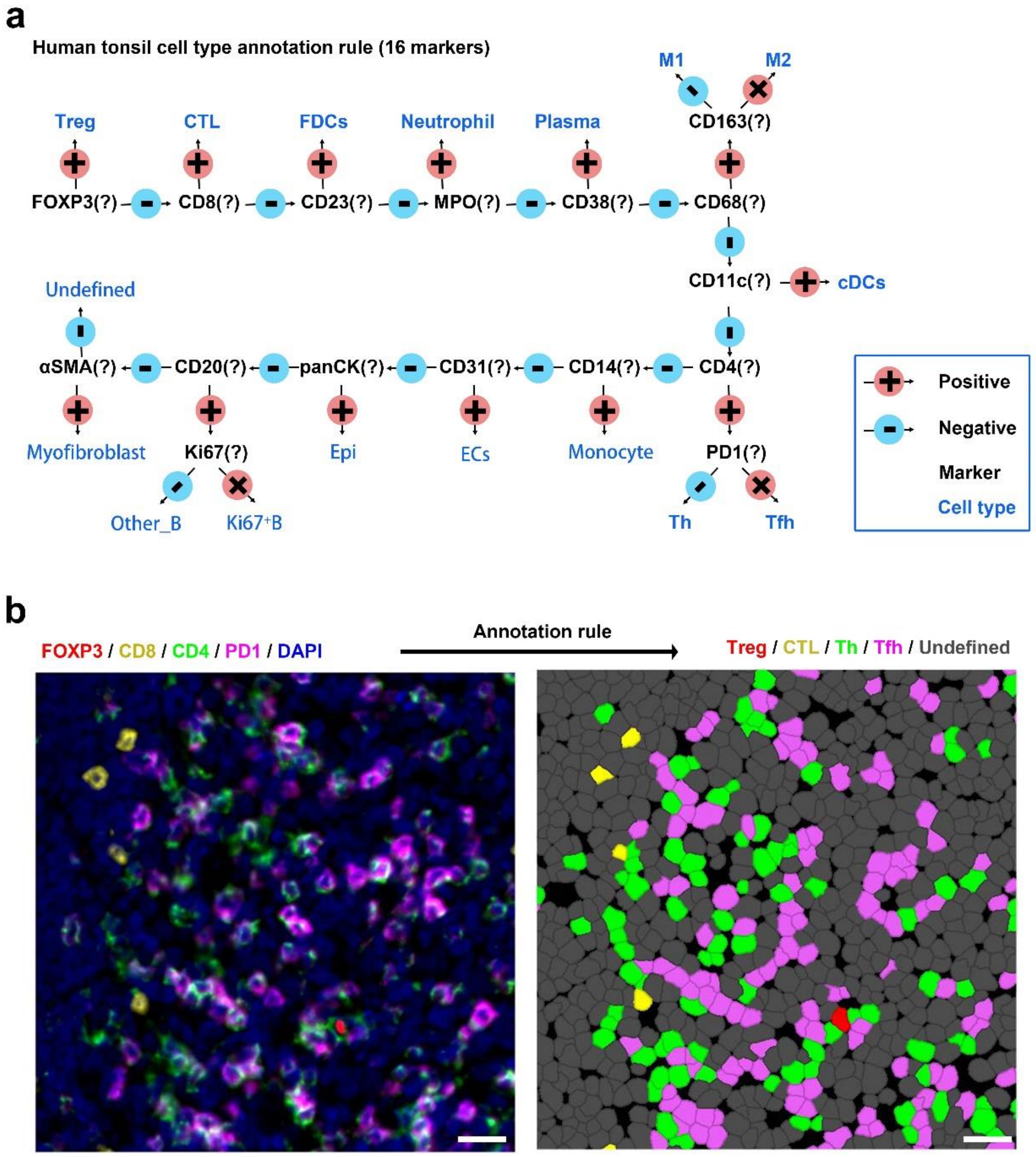


**Supplementary Fig. 5 Cell type annotation rule of the tonsil superplex image in fig5. D.**

(a) The cell type annotation rule based on typical lineage markers, expected population abundance, and staining quality supervision.

(b) In one of the ROIs in the Fig. 5d image, we used the specified annotation rule and obtained excellent ground-truth. Scale bar, 20 μm.

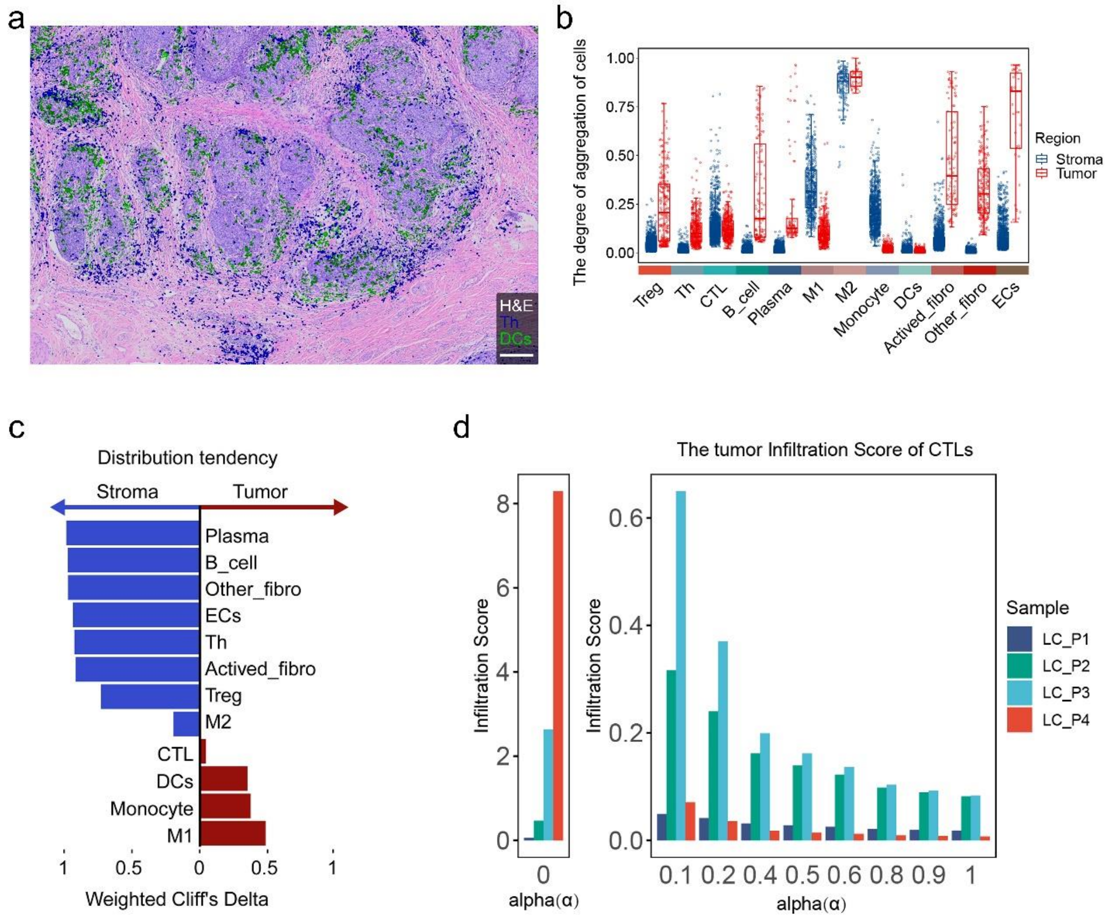


**Supplementary Fig. 6 Evaluation of spatial distribution tendencies for individual types of immune cells.**

(a) Th and DCs identified in Fig. 6a were overlay on the H&E image of the adjacent section, and the distribution tendency of different cells in the tissue could be seen. Scale bar, 200 μm.

(b) The degree of aggregation of various cells, excluding tumor cells, in the tumor and stroma areas. Values closer to 0 indicate stronger aggregation.

(c) The distribution tendency of different immune cells between stromal and tumor regions: Red bars represent cells that tend to accumulate in the stromal region, while blue bars represent cells that tend to accumulate in the tumor region. The larger the value, the stronger the tendency.

(d) The test results indicate that an alpha(α) value of 0.1 is the most suitable for calculating the tumor Infiltration Score of CTLs.

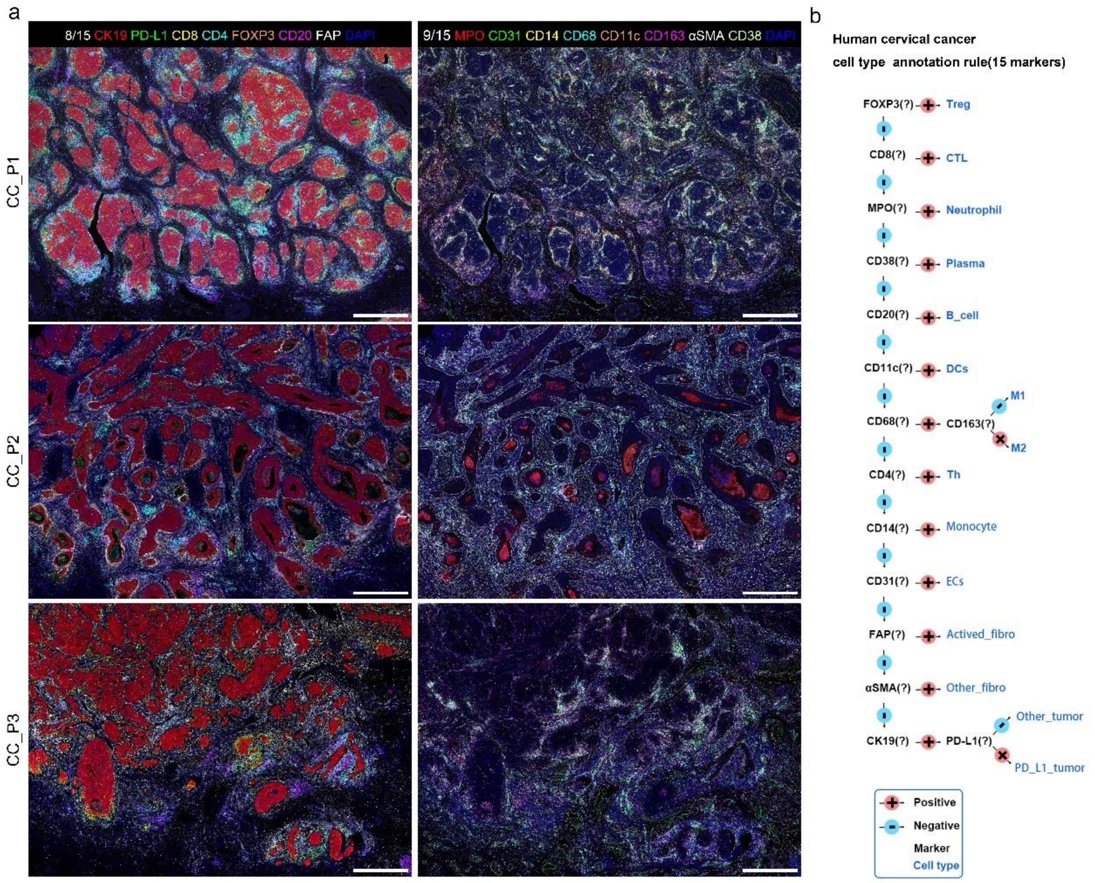


**Supplementary Fig. 7 The superplex raw images of three cervical cancer patients and the cell type annotation rule.**

(a) The superplex raw images of three cervical cancer patients. Scale bar, 1 mm.

(b) The cell type annotation rule of three cervical cancer patients.
